# Dendritic spine neck plasticity controls synaptic expression of long-term potentiation

**DOI:** 10.1101/2023.01.27.525952

**Authors:** Rahul Gupta, Cian O’Donnell

## Abstract

Dendritic spines host glutamatergic excitatory synapses and compartmentalize biochemical signalling underlying synaptic plasticity. The narrow spine neck that connects the spine head with its parent dendrite is the crucial structural element of this compartmentalization. Both neck morphology and its molecular composition differentially regulate exchange of molecular signals between the spine and rest of the neuron. Although these spine neck properties themselves show activity-dependent plasticity, it remains unclear what functional role spine neck plasticity plays in synaptic plasticity expression. To address this, we built a data-constrained biophysical computational model of AMPA receptor (AMPAR) trafficking and intracellular signalling involving Ca^2+^/calmodulin-dependent kinase II (CaMKII) and the phosphatase calcineurin in hippocampal CA1 neurons, which provides new mechanistic insights into spatiotemporal AMPAR dynamics during long-term potentiation (LTP). Using the model, we tested how plasticity of neck morphology and of neck septin7 barrier, which specifically restricts membrane protein diffusion, affect LTP. We found that spine neck properties control LTP by regulating the balance between AMPAR and calcineurin escape from the spine. Neck plasticity that increases spine-dendrite coupling reduces LTP by allowing more AMPA receptors to diffuse away from the synapse. Surprisingly, neck plasticity that decreases spine-dendrite coupling can also reduce LTP by trapping calcineurin, which dephosphorylates AMPARs. Further simulations showed that the precise timescale of neck plasticity, relative to AMPAR and enzyme diffusion and phosphorylation dynamics, critically regulates LTP. These results suggest a new mechanistic and experimentally-testable theory for how spine neck plasticity regulates synaptic plasticity.

## Introduction

Learning and memory in the brain is primarily mediated by synaptic plasticity: changes to the strength of the synaptic connections between neurons [1,2]. Excitatory glutamatergic synapses are typically located on dendritic spines, tiny protrusions from the dendritic shafts of a variety of neurons [3-5]. Synaptic strength correlates with dendritic spine head size [3,6-8]. Dendritic spines are highly compartmentalized structures in terms of electrical as well as chemical communications with the rest of the neuron [9-12]. They are, to a fair extent, endowed with their own machineries and rules for electrophysiological and biochemical computations, which critically shape the synaptic plasticity. The physical size and shape of dendritic spines affect the dynamics of their biochemical signals [13-16], but how these physical properties determine the functional ‘rules of plasticity’ remains unclear.

The specialized structure which is pivotal to the dendritic spine compartmentalization is the spine neck. It restricts the exchange of membrane as well as cytosolic chemical species between the spine head and the dendritic shaft [17-20]. Morphologically, spine necks are roughly cylindrical structures with typically narrow widths (0.1 to 0.3 *μm* diameter, [7,21-23]) and, thus, offer high resistance to the passage of ions and molecules. Anatomical studies of dendritic spines have revealed a wide distribution of spine neck morphology, in terms of its length and radius [7,21-23]. These studies have also showed lack of correlation between the spine neck morphology and the spine head size. Spine necks are also equipped with specialized cytoarchitectural molecules which can differentially regulate the flow of membrane and intracellular chemical species to a great distinction [24-26]. For instance, complex septin7 rings typically spotted at the base of the spine neck (close to the dendritic shaft) only hinder diffusion of membrane proteins in-and-out of the dendritic spines, leaving the intracellular cytosolic exchange entirely unaffected [24]. Akin to neck morphology, septin7 complexes also exhibit a wide range of distribution, with little to high degree of presence at the neck base [24].

Remarkably, spine necks are also highly dynamic structures and have been observed undergoing significant morphological changes during events of synaptic plasticity [26-30]. Often referred to as spine neck plasticity, it has been noted to critically modulate the electrical coupling [27,28] as well as Ca^2+^ exchange [31] between the dendritic spines and the shafts. Although changes in the spine neck morphology immediately imply rearrangements of the supporting cytoarchitectural molecules, correlation between the neck morphology and distributions of unique constituent molecules, such as septin7 complexes, has yet not been established. Therefore, it is unclear whether an identical underlying signalling process proportionately scale all the cytoarchitectural elements together up or down during changes in neck morphologies. If there exists a differential signalling regulation, it will broaden the influence of spine neck plasticity in terms of its morphological restriction and the additional capacity to differentiate between membrane and cytosolic exchanges.

Given that spine neck properties affect synapse biochemical signalling, it seems likely that spine neck plasticity affects synaptic plasticity. However, this aspect has yet remained unaddressed in the previous experimental as well as theoretical studies. The possibility of the modulation of synaptic strengths through morphological changes in the spine neck has been hypothesised long ago [32]. In the study from Araya et al. [27], shrinkage of spine neck length in response to spike timing-dependent plasticity protocols were shown to enhance the contributions of dendritic spine stimulations to the evoked somatic potential. Tonnesen et al. [28] also demonstrated a similar observation where long-term potentiation (LTP) of stimulated spines led to wider and shorter neck. However, in all these studies, a clear distinction between changes in the synaptic properties and electrotonic attenuation through spine neck could not be established. Previous computational modelling studies have also extensively explored intracellular signalling and AMPA-type receptor (AMPAR) trafficking during events of synaptic plasticity [33-40]. However, the effect of simultaneous spine neck plasticity on the LTP expression is missing.

In this study, we addressed this issue by building and studying a data-constrained biophysical model of AMPAR trafficking, phosphorylation and the major upstream enzymatic signalling, which involved Ca^2+^/Calmodulin-dependent kinase II (CaMKII) and the phosphatase calcineurin. CaMKII action is necessary for hippocampal LTP, while calcineurin action is necessary for LTD [13,15]. Even though our focus was on single-synapse plasticity, to accurately capture the spatial dynamics we modelled an entire apical dendritic branch of a CA1 hippocampal pyramidal neuron with hundreds of dendritic spines. To constrain the model parameters, we fit to fluorescence imaging data from Patterson et al. (2010) [41] on AMPAR trafficking in CA1 apical dendritic branches during the resting unstimulated condition, as well as during single spine stimulation for long-term potentiation (LTP). In addition to the structural plasticity of the stimulated spine head (spine head enlargement) reported by Patterson et al. [41], we also implemented spine neck plasticity in terms of changes in the neck morphology as well as the neck septin7 barrier. We considered the possibility of independent changes in both the neck properties to occur at different timescales. The present findings suggested that spine neck plasticity could dramatically influence the expression of LTP by modulating the leakage of LTP-associated signals from the stimulated spine head.

## Results

We designed a reaction-diffusion model for the dynamics of AMPAR trafficking and intracellular CaMKII-Calcineurin signalling in a thin apical 100 *μm* long [42] dendritic branch (Figure 1A) of a typical CA1 pyramidal neuron (see Methods). 100 identical mushroom-shaped spines were located on a uniformly thick (*r*_*dend*_ = 0.5 *μm*) dendritic shaft [42] with a regular inter-spine distance of 1 *μm* [43]. Each spine (Figure 1B) had a cylindrical spine neck (*r*_*neck*_ = 0.0562 *μm, l*_*neck*_ = 0.3 *μm*) [7,21] and a spherical spine head (*A*_*spine*_ = 1.1 *μm*^2^, *V*_*spine*_ = 0.1 *μm*^3^) [7,21,43]. The membrane of the spine head was divided into a small postsynaptic density (PSD) region (*A*_*PSD*_ = 0.1 *μm*^2^) and the remaining extrasynaptic membrane region (ESM, *A*_*ESM*_ = 1 *μm*^2^) [21,43,44]. The dendritic shaft was also segmented into thin 0.1 *μm* long compartments along the length of the shaft, for the purpose of simulation with high spatiotemporal precision. The modelled dendritic branch was connected at its origin to the parent primary dendritic shaft, which acted as a steady source of AMPARs, and CaMKII and calcineurin enzymes.

**Figure 1.**
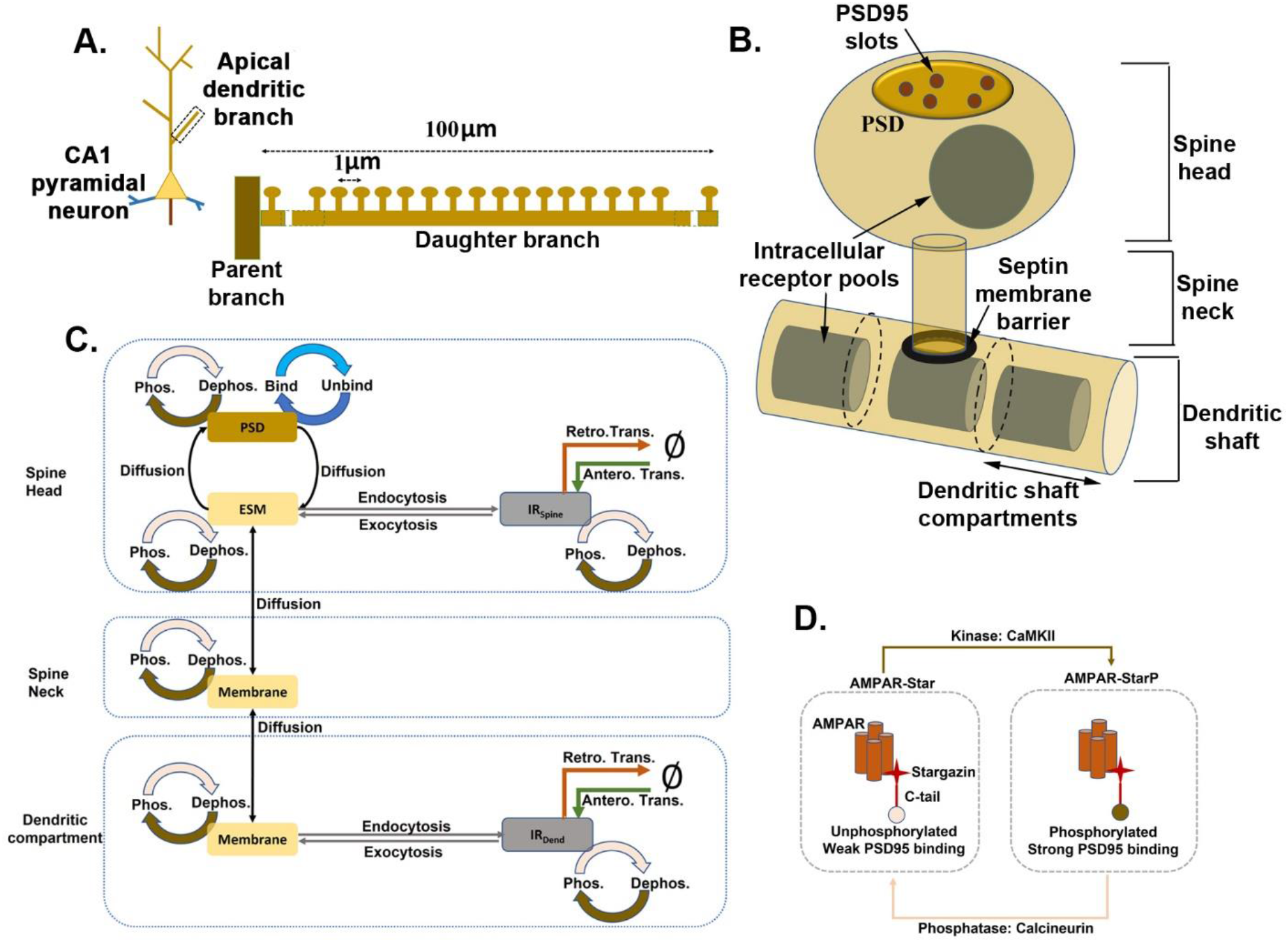
Schematic description of the model. (A) The model consisted of the dynamics of AMPAR trafficking and CaMKII-calcineurin intracellular signalling across a 100 *μm* long apical dendritic branch of a CA1 pyramidal neuron, with hundreds of dendritic spines located at a regular distance of 1 *μm*. (B) A dendritic spine had a head and a neck. Postsynaptic density (PSD) located in the top membrane region of the spine head, facing presynaptic glutamate release zone, contained AMPAR-binding PSD95 proteins. Septin ring located at the base of the spine neck restricted diffusion of only membrane proteins in-and-out of the spine. The dendritic shaft was uniformly divided into 0.1 *μm* long shaft compartments. Intracellular pools of AMPARs were located in the spine head as well as dendritic shaft compartments. (C) Various components of the AMPAR trafficking and receptor phospho-/dephosphorylation dynamics are shown. (D) The CaMKII-dependent phosphorylation of the cytosolic C-tails of the stargazin proteins, constantly associated with AMPARs, enhanced receptor binding with the PSD95 proteins. Calcineurin-dependent dephosphorylation of AMPAR-stargazin complex weakened the binding.

The model framework had two concurrently functioning modules: AMPAR trafficking and CaMKII-calcineurin intracellular signalling (Figure 1C). AMPARs in the model were assumed to always exist as AMPAR-stargazin complex [45]. Stargazin is a member of transmembrane AMPAR regulatory proteins (TARPs) and is required by the AMPARs to bind with the PSD95 scaffolding proteins in the synapse [45,46]. AMPAR trafficking involved: (a) exchange of AMPARs amongst membrane compartments through lateral diffusion in the membrane [19,47], (b) exocytosis and endocytosis of AMPARs between the membrane and intracellular pool [48,49], (c) loss and gain of AMPARs in the intracellular pools through synthesis/degradation [39,50] and retrograde/anterograde vesicular transports [51,52] and (d) binding and unbinding of AMPARs with the PSD95 slot proteins [45,46]. In the intracellular signalling module, active CaMKII and calcineurin in a compartment locally drove phosphorylation and dephosphorylation of AMPAR-stargazin complexes, respectively [53,54]. Phosphorylated AMPAR-stargazin complexes bound with the synaptic PSD95 slot proteins with a stronger affinity [53,54] than the dephosphorylated species (Figure 1D). In our model, this differential binding of the two AMPAR-stargazin species was the core mechanism underlying LTP expression in terms of change in the synaptic AMPAR count within the PSD region of the stimulated spine [53,54].

### Baseline Configuration of the Model

The baseline configuration of the model represented the initial unstimulated steady-state condition of the AMPAR trafficking and the intracellular signalling in the dendritic branch. We set the baseline model’s parameter values for spine morphology, receptor numbers, PSD95 slot proteins and the enzymes (CaMKII and calcineurin) to their empirical estimates from the literature (Methods). AMPARs were present at a uniform density of 10 *μm*^−2^ in the dendritic shaft membrane as well as the spine head ESM and the neck membrane [55,56]. Each spine head and 1 *μm* length of the dendritic shaft contained an intracellular pool of 100 AMPARs [48,49]. There were 100 AMPAR-binding PSD95 slot proteins in the PSD [46,57]. Only 25% of the slot proteins were occupied with AMPARs, in accordance with the empirically observed abundance of PSD95 proteins unbound with AMPARs in the unstimulated conditions [46,57,58]. Owing to the phospho-dephosphorylation dynamics of stargazin, we had two subpopulations of AMPAR-stargazin complexes: unphosphorylated (AMPARstar) and phosphorylated (AMPARstarP) complexes. 90% of the total AMPARs in the membrane as well as intracellular pools were AMPARstar in the baseline condition. The ratio was reversed for the PSD95-bound AMPARs, as the AMPARstarP bound strongly with the PSD95 proteins.

Because we were concerned with AMPAR surface expression dynamics on a timescale of seconds to minutes, we did not explicitly model the complex millisecond-timescale dynamics of NMDA receptor-dependent Ca^2+^-mediated activation of CaMKII and calcineurin during glutamatergic stimulation [35,39,59]. Instead, we directly varied the concentrations of active CaMKII and calcineurin in the stimulated spine head based on their empirically observed temporal waveforms [35,39,60,61]. We implicitly modelled CaMKII activity by varying phosphorylation rate constant in the stimulated spine head, as experiments have found that activated CaMKII remains confined within the stimulated spine [60-63]. In contrast, calcineurin is known to diffuse out from stimulated spines [14,62]. Therefore, we modelled calcineurin explicitly as a cytosolic enzyme which diffuses between spines and the dendrite. We assumed a homogeneous distribution of the CaMKII enzymes, and hence the corresponding baseline phosphorylation rate constant, and the calcineurin enzymes across the entire unstimulated dendritic branch (Methods).

Next, we set the baseline rate parameters of AMPAR trafficking processes. AMPARs constantly exhibit lateral diffusion in the membrane between dendritic spines and shaft [19,47]. The empirical estimates of the diffusion coefficient of AMPARs in the obstacle-less ESM of the spine head (*D*_*ESM*_) and dendritic shaft (*D*_*dend*_) is 0.1 *μm*^2^. *s*^−1^ [19,47]. However, due to the presence of immense molecular crowding in the PSD, the local diffusion coefficient (*D*_*PSD*_) is empirically estimated to be roughly one-tenth of that in obstacle-less membrane [47,64-66]. Further, presence of cytoarchitectural proteins such as septin7 barriers in the spine neck reduces the receptor diffusion across the neck [24], which is experimentally estimated to be *D*_*neck*_ = 6.7 × 10^−3^ *μm*^2^. *s*^−1^ [19]. Therefore, we considered that all the spine necks in the baseline configuration have identical septin7 barrier ring located at their bases (Figure 1B) and an identical *D*_*neck*_. Using these empirical estimates and the geometry of the membrane compartments, we computed the rate constants of receptor hopping amongst the compartments (Methods).

AMPAR trafficking also involved exchange of receptors between the membrane and the IR pools through exo- and endo-cytosis in the spine head as well as the dendritic shaft compartments. The IR pools also gained and lost receptors through synthesis/degradation as well as motor protein-based active transport. We estimated the baseline values of these parameters by using automatic optimization methods to fit the model to the fluorescence imaging data from Patterson et al. (2010) [41] on AMPAR trafficking under unstimulated conditions (Methods). This fluorescence-recovery after photobleaching (FRAP) study monitored the fluorescence-recovery of SEP-tagged GluA2-containing AMPARs in the spine and dendritic shaft after photobleaching of a stretch of dendrite. The model fitted well to the experimental data (Figure 2A). After establishing the initial baseline configuration of the model, we proceeded to simulate the conditions of spine stimulation.

**Figure 2.**
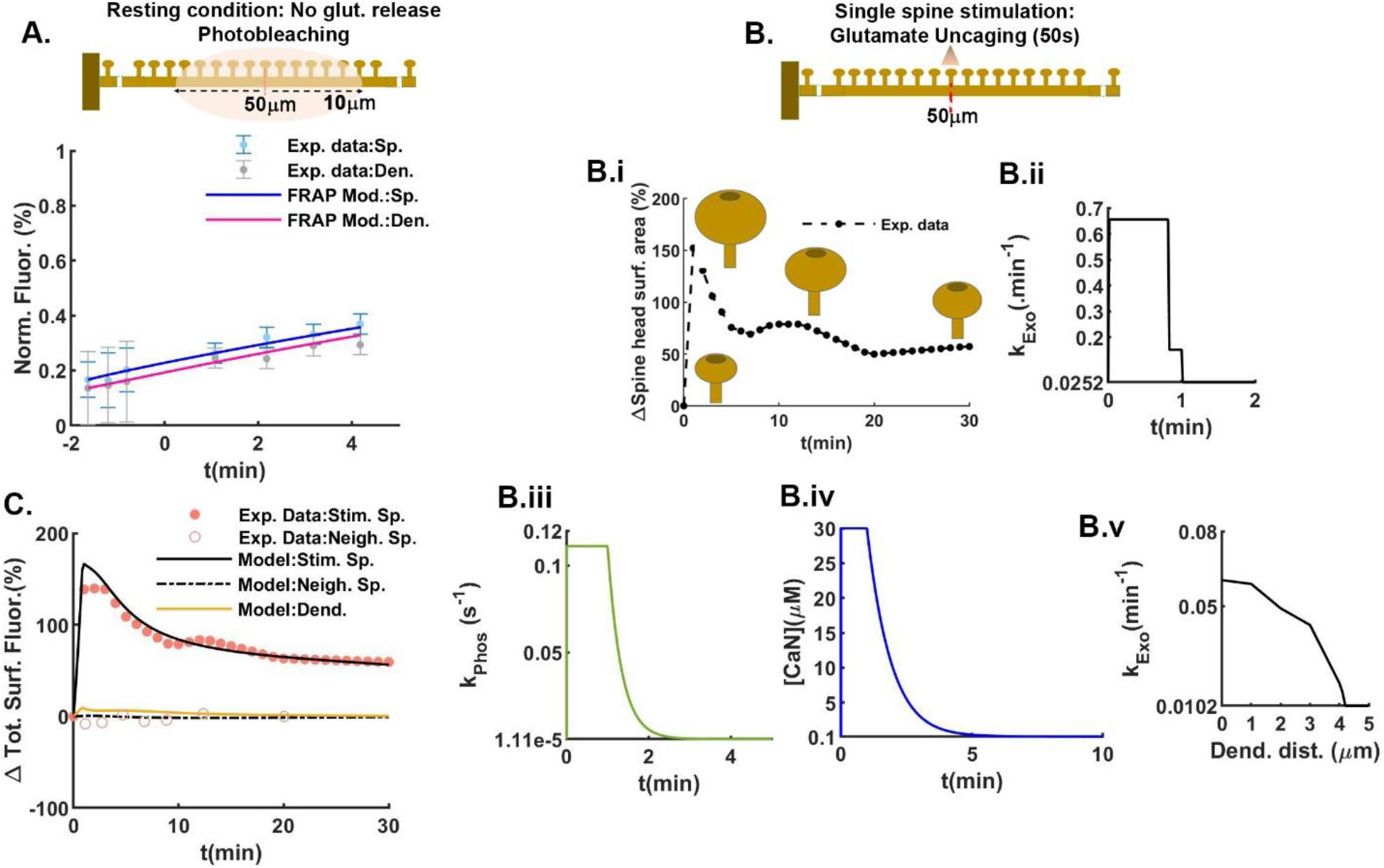
Validating the Model through reproducing experimental data. The fluorescence imaging data from the Patterson et al. (2010) study [41] on AMPAR trafficking in a proximal apical dendritic branch of a CA1 pyramidal neuron was used to validate the model. (A) In the resting unstimulated condition of the dendritic branch, the model closely reproduced the experimentally-observed recovery of surface AMPAR fluorescence in the dendritic spine heads and shaft, after the photobleaching of 20 *μm* stretch around the centre of the dendritic branch. The fluorescence recovery was normalized to the baseline surface AMPAR fluorescence before the photobleaching. (B) Glutamatergic stimulation (via. a minute long glutamate uncaging protocol) of a single spine located at the centre of the dendritic branch was implemented in silico through the direct empirically-guided temporal changes in the spine head size (B.i, except PSD), AMPAR exocytosis rate constant (B.ii), the CaMKII-dependent AMPAR phosphorylation rate constant (B.iii), and the concentration of activated calcineurin phosphatase (B.iv), within the stimulated spine head. The exocytosis rate constant in the dendritic shaft underneath the stimulated spine was also step increased during glutamatergic stimulation, similar to that in the stimulated spine. (B.v) However, the intensity of the step-increase in the dendritic exocytosis rate constant gradually decreased away from the location of the stimulated spine. (C) The model closely reproduced the experimental data on the change in the total surface AMPAR fluorescence in the stimulated spine head and the unstimulated neighbouring spine head for the single spine stimulation. ‘Total’ refers to the AMPAR population in the spine head ESM as well as PSD. AMPAR fluorescence is proportional to the AMPAR counts.

### AMPAR dynamics during single spine stimulation

To replicate the single spine glutamate-uncaging experiments by Patterson et al. [41], we chose the middle-most spine (located at 50 *μm* from the branch origin, Figure 2B) in our model as the target of glutamatergic spine stimulation. We implemented the LTP-inducing spine stimulation in three parts, briefly (see Methods for details): (1) We directly increased the size of the spine head (Figure 2B.i), in a manner identical to the temporal profile of spine head enlargement observed by Patterson et al. [41]. The size of the PSD and the number of PSD95 slot proteins were not altered [67,68]. (2) We step-increased the exocytosis rates of the AMPAR species in the stimulated spine as well as the dendritic shaft (Figures 2B.ii and 2B.v), again in a spatiotemporal manner as reported by Patterson et al. The endocytosis rate constants, and the AMPAR dynamics within the intracellular pools, were unaffected. (3) At the instance of glutamate uncaging, we step-increased and clamped the phosphorylation rate constant (Figure 2B.iii) and the concentration of active calcineurin (Figure 2B.iv) for the entire 50 seconds duration of glutamate uncaging [35,39,60,61]. Following the stimulation, the phosphorylation rate constant and the concentration of active calcineurin were allowed to exponentially decay to their baseline values. Also, the active calcineurin intracellularly diffused from the stimulated spine into the dendritic shaft. The values for the magnitude of step-increases, decay time-constants, and the diffusion coefficient of calcineurin were estimated in order to reproduce the temporal profile of total surface AMPAR fluorescence in the stimulated spine head reported by Patterson et al. [41].

In order to dissociate the relative contributions of enhanced AMPAR exocytosis versus intracellular signalling during spine stimulation to the dynamics of AMPAR surface expression, we performed perturbation simulations where each of the two processes was removed in turn (Figure 3A). These simulations showed that enhanced exocytosis dominates for the initial 2-3 minutes post-stimulation (light blue curve, Figure 3A), but signalling dominates from 5 minutes post-stimulation, after exocytosis has returned to baseline (dark blue curve, Figure 3A). Note that this switch at 5-minutes occurred later than the CaMKII activity transient, which lasted only ∼2 minutes (Figure 2B.iii). The delay was due to the fact that phosphorylated AMPAR-stargazin complexes take time to gradually bind to the PSD95 slots and accumulate at the synapse. These results support the established idea that both enhanced AMPAR exocytosis and intracellular signalling during spine stimulation are important for LTP expression, but at different temporal phases [69].

**Figure 3.**
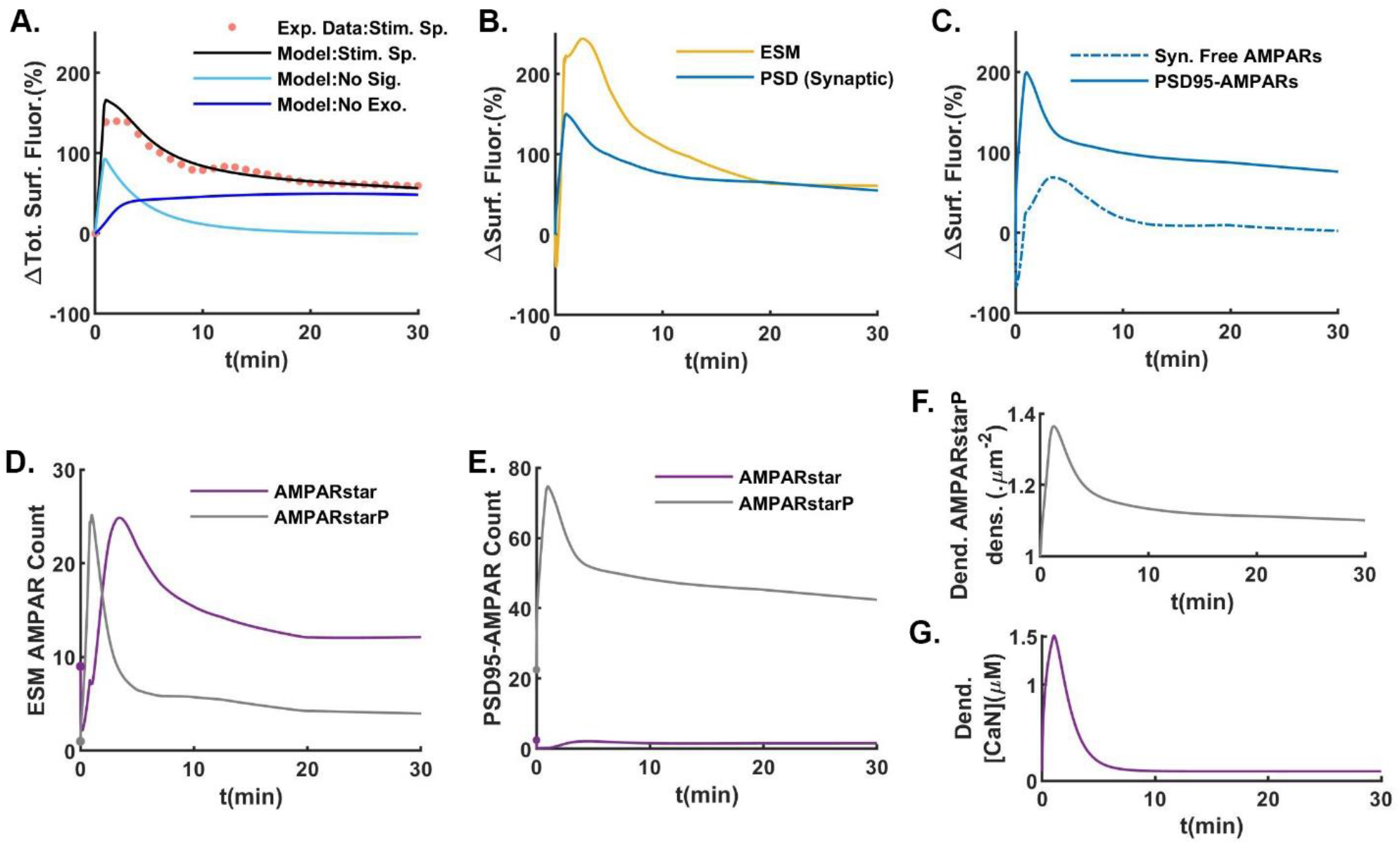
Dynamics of surface AMPARs under single spine stimulation. (A) Simultaneous changes in the kinetics of the AMPAR exocytosis as well as of the intracellular CaMKII- and calcineurin-mediated AMPAR phosphorylation and dephosphorylation, respectively, during spine stimulation were required to reproduce the empirical profile of change in the total surface AMPAR fluorescence in the stimulated spine head. Surface AMPAR fluorescence resulting from the absence of either of the processes are shown to highlight their individual contributions. (B) The change in total surface AMPAR fluorescence in the stimulated spine head consisted of the changes in the PSD and ESM regions of the spine head. The surface AMPAR fluorescence in the PSD accounted for synaptic AMPAR count and, hence, represented potentiation of the synaptic strength (LTP). (C) Both freely diffusing and PSD95-bound AMPARs in the PSD contributed to the synaptic surface AMPAR fluorescence. (D) Dynamics of phosphorylation and dephosphorylation of AMPAR-stargazin complex in the stimulated spine head are shown through the temporal profiles of the surface expression of unphosphorylated (AMPARstar) and phosphorylated (AMPARstarP) species in the ESM region of the stimulated spine head. (E) The same is shown for PSD95-bound species in the PSD region of the stimulated spine head. (F) and (G) show leakage of the AMPARstarP and active calcineurin phosphatase, respectively, into the dendritic shaft just underneath the stimulated spine head.

To ask how the AMPAR population differed in the ESM versus the PSD of the stimulated spine head, we plotted the dynamics of both populations during the same simulation as above (Figure 3B). While the AMPAR population in the PSD rapidly rose at the instance of stimulation, the ESM population initially showed a sharp decline immediately followed by a rapid rise <1 minute post-stimulation. Within the PSD, the population of PSD95-bound AMPARs rose whereas the freely diffusing AMPAR population initially dropped before recovering to a net positive change 1-2 minutes post-stimulation (Figure 3C). In the ESM, AMPARs were rapidly phosphorylated causing an initial switch from AMPARstar dominating to AMPARstarP dominating (Figure 3D). The phosphorylated fraction of PSD95-bound AMPARs also rapidly increased (Figure 3E).

Together, these results support the following description for AMPAR dynamics during LTP: initially in the first minute, during stimulation, there is a decline in the total receptor population in the ESM due to phosphorylated AMPARs diffusing from the ESM into the PSD and binding to the abundant free PSD95 slot proteins. Then, after stimulation ends, CaMKII activity returns to baseline within 2 minutes post-stimulation, but active calcineurin took longer to decay, ∼5 minutes. In this 2-5 minutes period where phosphatase activity dominates, the AMPARstarP population in both the ESM (grey curve, Figure 3D) and PSD (grey curve, Figure 3E) becomes rapidly dephosphorylated. The dephosphorylation of PSD95-bound AMPARs causes them to unbind and diffuse freely. This causes a bump in free AMPARs in the PSD at 5 minutes post-stimulation (dashed curve, Figure 3C). Meanwhile, a portion of both AMPARstarP (Figure 3F) and calcineurin diffused out from the stimulated spine to the dendritic shaft (Figure 3G). From 5-30 minutes post-stimulation, the receptor populations across all the membrane compartments gradually decreased via the baseline AMPAR trafficking mechanisms.

### Effects of spine neck plasticity on LTP dynamics

Next, we explored how spine neck plasticity at the stimulated spine (Figure 4A) could affect the LTP expression. We implemented both changes in the spine neck morphology (neck morphological plasticity) as well as alterations in the septin7 barrier (neck septin plasticity). Since it is not known if they are linked, for simplicity we assumed they occur independently. Biologically, both neck radius and length affect the degree of coupling between the spine and dendritic shaft [14,17,20,62]: decrease in neck length and increase in neck radius increase the spine-dendrite coupling, for both surface and intracellular diffusion, and vice versa (Figure 4B). Based on the empirical distributions of the spine neck radii and lengths [7,21,43], we considered two extreme situations of neck restrictions: (a) widened-and-shortened neck with minimum neck restriction (*r*_*Neck*_ = 0.1262 *μm, l*_*Neck*_ = 0.1250 *μm*) and (b) narrowed-and-elongated neck with maximum neck restriction (*r*_*Neck*_ = 0.03 *μm, l*_*Neck*_ = 1 *μm*). All other possible combinations of neck radii and lengths give a spectrum of neck restrictions bounded by these two extreme conditions (Figure 4B). Accordingly, we called the baseline neck morphology used in the above simulations the ‘original’ neck, because it had the most common morphology values [7,21,43]. For the neck septin plasticity, we either had intact baseline septin barrier, referred to as the septin-On state, or lack of septin barrier, referred to as the septin-Off state. Therefore, combining the independent alterations of neck morphology and septin7 barrier, we had six possible states of spine neck at the stimulated spine (Figure 4A).

**Figure 4.**
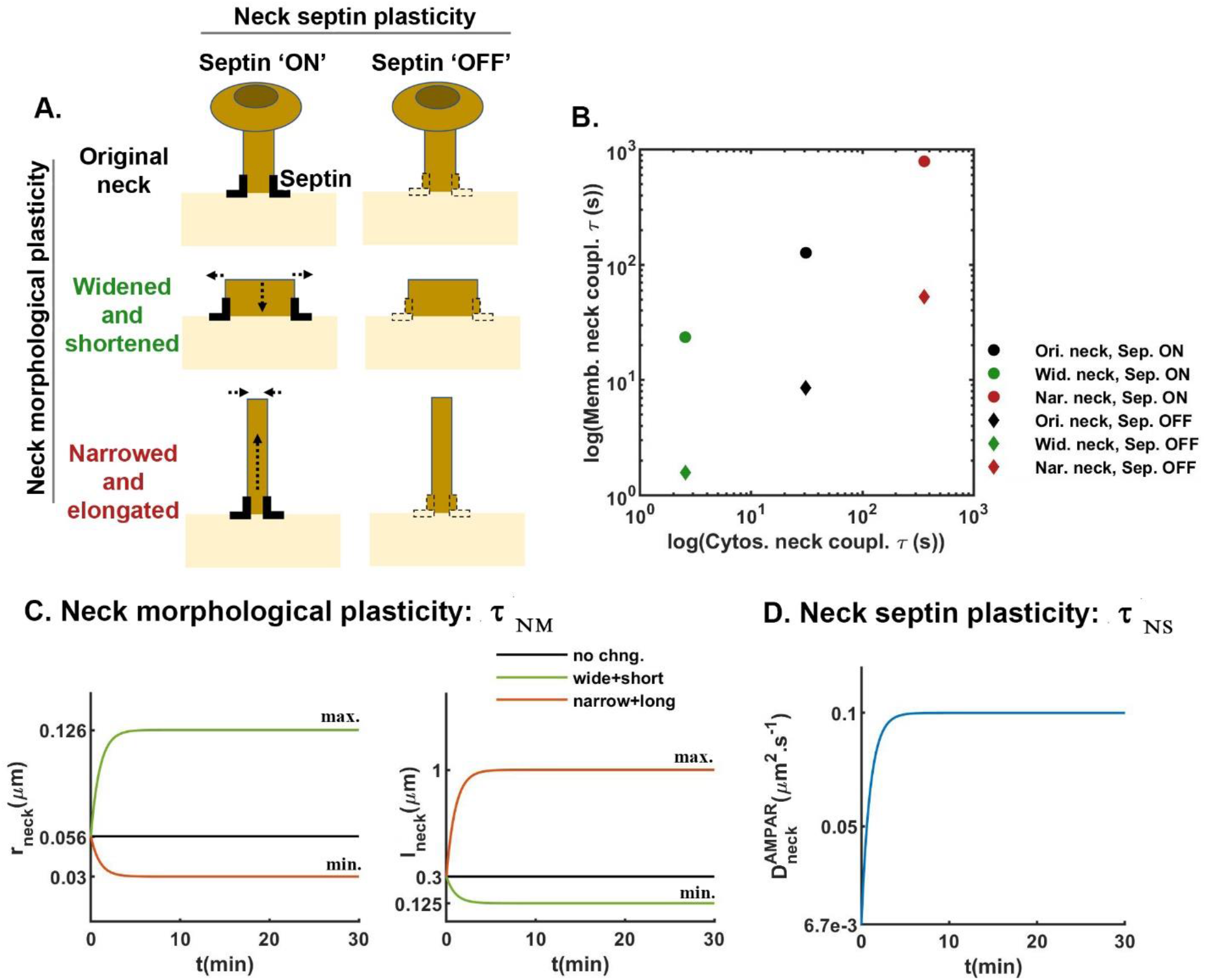
Schematic description of spine neck plasticity. (A) Neck of the stimulated spine could undergo morphological plasticity as well as septin plasticity, independently. Besides the initial baseline neck morphology (referred to as the original neck), two other extremes of neck morphologies pertaining to the maximum (widened-and-shortened neck) and minimum (narrowed-and-elongated) conductance of the spine neck were considered. Spine neck with intact septin7 barrier represented septin-On state. Neck septin plasticity involved complete fading of the septin7 barrier, referred to as the septin-Off state. (B) The coupling strengths between spine head and dendritic shaft for the membrane diffusing AMPAR and cytosol diffusing calcineurin signals under the different neck morphological and septin states are shown in terms of diffusion time constants. (C) Dynamic neck morphological plasticity involved simultaneous changes in the neck radius and length with a time constant, *τ*_*NM*_. For the plasticity towards widened-and-shortened neck, the neck radius and length exponentially increased and decreased to their empirically-known maximum and minimum values, respectively (shown in green). The reverse occurred for the plasticity towards narrowed-and-elongated neck (shown in red). (D) Neck septin barrier affected AMPAR diffusion constant in the neck. Therefore, neck septin plasticity essentially signified rise in the neck AMPAR diffusion coefficient from the initial very low value to the maximum value of AMPAR diffusion coefficient in the free dendritic membrane. Septin plasticity occurred exponentially with a time constant, *τ*_*NS*_.

Since the timescales of neck morphological plasticity are poorly quantified, we defined a neck plasticity time constant parameter *τ*_*NM*_. After induction of spine neck plasticity, neck radius and length simultaneously and exponentially varied at the *τ*_*NM*_ timescale (Figure 4C). Similarly for the neck septin plasticity, we defined a time constant *τ*_*NS*_ with which the AMPAR membrane diffusion coefficient in the neck exponentially increased to the prescribed maximum value (Figure 4D) of AMPAR diffusion coefficient in the free dendritic membrane.

We then reran the same single-spine stimulation experiment as in Figure 3, with added spine neck morphological and septin plasticity. Overall, we observed a strong impact of spine neck plasticity on the expression of LTP (Figure 5). Since both neck morphological and septin plasticity could occur with different values of *τ*_*NM*_ and *τ*_*NS*_, respectively, we began with the case of extremely fast or instantaneous neck plasticity events by setting both time constants to 1 second. Although neck morphological plasticity towards either the widened-and-shortened (Figure 5A.i) or the narrowed-and-elongated (Figure 5A.iii) neck configurations did not affect the initial transient of LTP, it did lead to faster subsequent decay in the total spine head surface AMPAR count (total fluorescence) post-stimulation compared to the original neck (Figure 5Aii.). The decay was fastest and deepest for the narrowed-and-elongated neck case. Concurrent neck septin plasticity increased the decay rate for all neck morphologies.

**Figure 5.**
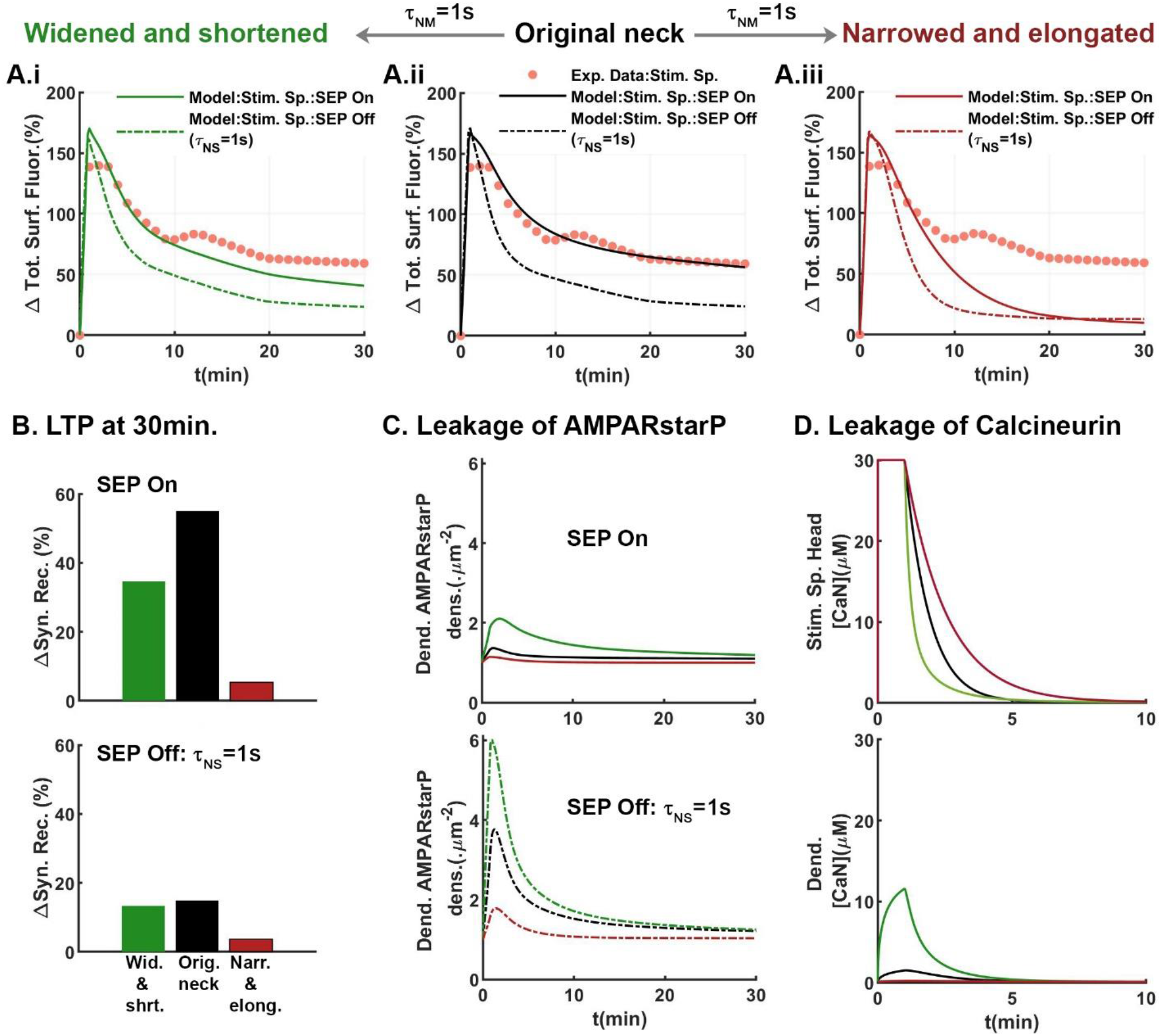
Spine neck plasticity strongly affects LTP expression at the stimulated spine. (A) The temporal profiles of change in the total surface AMPAR fluorescence in the stimulated spine head under the different spine neck plasticity are shown, along with the experimental imaging data. Spine neck morphological plasticity occurred instantaneously with the time constant *τ*_*NM*_ = 1 second and spine neck septin plasticity occurred with *τ*_*NS*_ = 1 second. (B) Magnitude of LTP at the stimulated spine in terms of change in the synaptic receptor count remaining by 30 minutes post-stimulation is shown under different spine neck plasticity. (C) Leakage of AMPARstarP into the dendrite shaft just underneath the stimulated spine is shown as rise in the local dendritic AMPARstarP membrane density. (D) Calcineurin also leaked from the stimulated spine head into the dendritic shaft, depending on the spine neck morphological plasticity. All the data for the original, the widened-and-shortened, and the narrowed-and-elongated necks are consistently shown in black, green and red colours, respectively.

We quantified the amplitude of sustained LTP as the change in the synaptic receptor count at 30 minutes post-stimulation. This strongly varied depending on spine neck plasticity (Figure 5B). The most prominent result was that plasticity that weakened the septin7 barrier at the base of spine neck substantially reduced the magnitude of LTP. The second key result was that neck morphological plasticity towards the narrowed- and-elongated neck configuration reduced LTP expression, regardless of the neck septin state. Neck widening-and-shortening also reduced LTP, but to a lesser extent.

The dynamics of the AMPAR surface population in the stimulated spine head was also simultaneously associated with the leakage of the two competing signals, AMPARstarP (Figure 5C) and calcineurin (Figure 5D), involved in local LTP expression. Spine neck coupling between the stimulated spine head and the dendritic shaft increased and decreased from neck morphological plasticity towards the widened-and-shortened and narrowed-and-elongated neck configurations, respectively. Accordingly, the leakage of both the signals systematically increased while going from the latter to the former neck morphologies, with the original neck showing intermediate intensity of leakage. Since leakage of AMPARstarP also depended on the neck septin7 barrier, neck plasticity that removes the septin barrier enhanced the leakage of AMPARstarP, with the precise magnitude determined by the neck morphological state (Figure 5C). This explained the abovementioned negative effect of neck septin plasticity on LTP expression at the stimulated spine head across all neck morphologies.

Leakage of calcineurin also impacted the decay profile of active calcineurin in the stimulated spine head (Figure 5D). Compared to the original neck morphology, stronger calcineurin leakage through the widened- and-shortened neck morphology led to faster decay of active calcineurin in the stimulated head, whereas the narrowed-and-elongated neck morphology prolonged the active calcineurin transient by trapping it in the spine head. This explained the observation that the narrowed-and-elongated neck configuration with very low spine neck coupling substantially reduced LTP magnitude (Figure 5B), as the prolonged presence of active calcineurin in the stimulated spine head strongly dephosphorylated AMPARs (Figure 5A.iii). Although the widened-and-shortened neck morphology decreased active calcineurin level in the stimulated spine head, which on its own should enhance LTP, the concurrent stronger leakage of AMPARstarP (Figure 5A.i) led to a net decrease in LTP magnitude (Figure 5B).

Next, we explored the effect of different timescales of the neck morphological and septin plasticity on LTP magnitude (Figure 6). Smaller values of *τ*_*NM*_and *τ*_*NS*_ led to faster neck morphological and septin plasticity, respectively. With intact septin barrier (no neck septin plasticity, Figure 6: bottom row), the LTP magnitude showed a non-monotonic dependence on the timescale of neck morphological plasticity. Very slow neck plasticity (*τ*_*NM*_ > 30 minutes) towards narrowed-and-elongated morphology (red bars on right side) caused an increase in LTP magnitude. However, further decreases in *τ*_*NM*_ led to gradual reduction in LTP magnitude, with a sharp decline around *τ*_*NM*_ = 1 minute. On the other hand, neck plasticity towards the widened-and-shortened morphology (green bars on left side) at slow timescales (*τ*_*NM*_ > 10 minutes) caused a reduction in LTP magnitude. Further decreases in tauNM caused a slight rise in the LTP magnitude, maximally around *τ*_*NM*_ > 1 minute, followed by a decreasing magnitude for extremely fast seconds-timescale neck plasticity. This non-monotonic variation of LTP magnitude under neck morphological plasticity remained largely conserved when neck septin plasticity also simultaneously occurred (Figure 6: top four rows). However, faster neck septin plasticity caused consistent decrease in the overall LTP magnitude.

**Figure 6.**
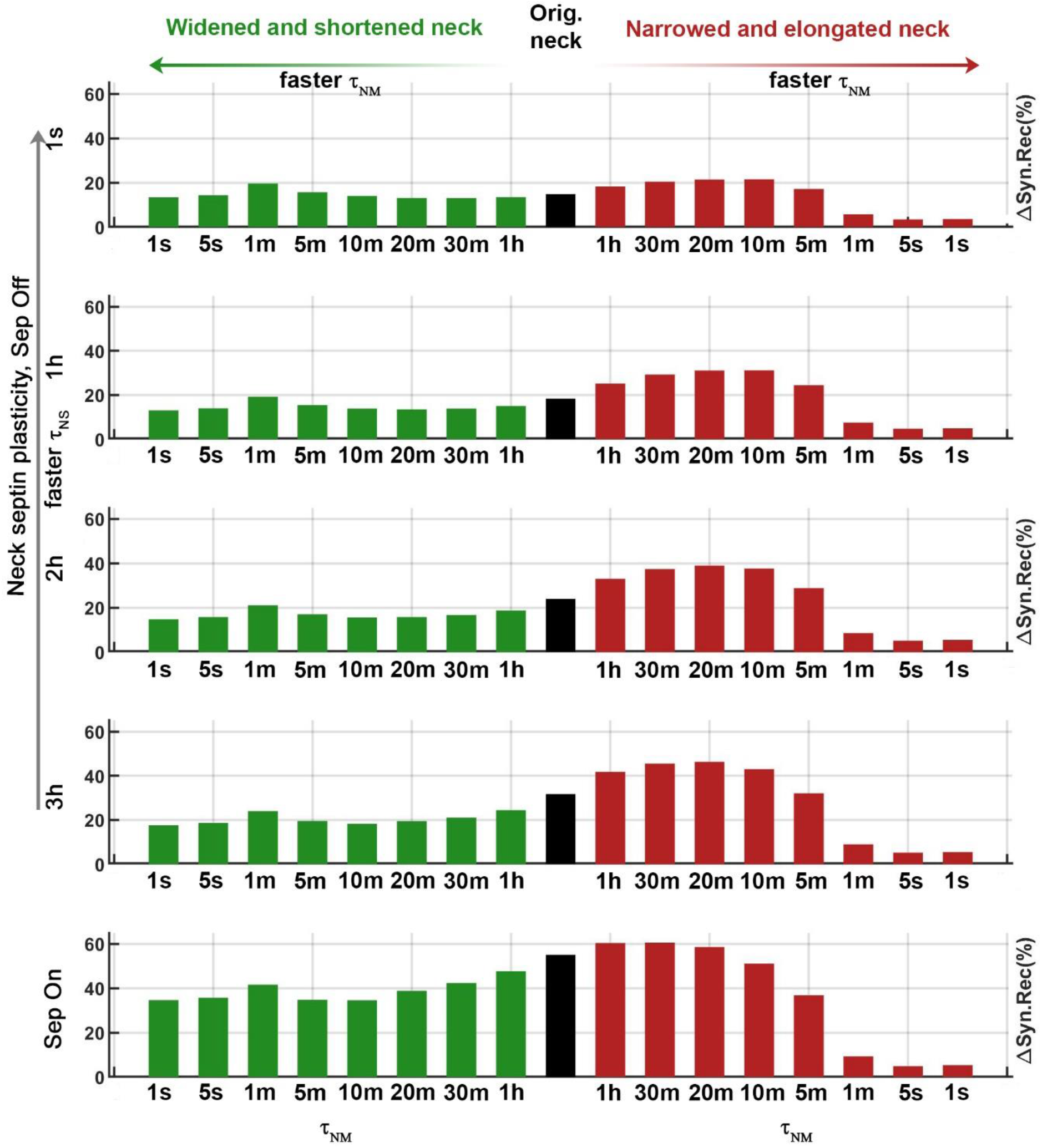
LTP magnitude is affected by the timescales of neck morphological and septin plasticity at the stimulated spine. LTP magnitude at the stimulated spine head is shown in terms of change in synaptic AMPAR count by 30 minutes post-stimulation. Rows of bar plots represent the neck septin plasticity. The bottom row represents absence of neck septin plasticity and intact septin barrier (Sep On state). The top four rows show neck septin plasticty causing removal of septin barrier (Sep Off states) with *τ*_*NS*_ decreasing from down to up. In each row, LTP magnitudes (bars) are shown for the two different directions of the neck morphological plasticity, the widened-and-shortened (green) and the narrowed-and-elongated neck (red) configurations, with the different timescales *τ*_*NM*_ at the stimulated spine head. LTP magnitude for the original neck is shown in black. ‘s’, ‘m’, and ‘h’ denote seconds, minutes, and hours, respectively.

Above we observed that the leakage dynamics of active calcineurin through the spine neck (Figure 5D, bottom panel) happened roughly within the first 5 minutes after spine stimulation, with maximum leakage occurring around 1 minute post-stimulation. AMPAR leakage also maximally occurred around 1-2 minutes post-stimulation but decayed over a longer span of 20-30 minutes (Figure 5C). Therefore, if the neck morphological plasticity is slow such that *τ*_*NM*_ is much larger than 5 minutes, it will largely affect the AMPARstarP leakage from the stimulated spine. However, as the morphological plasticity will get faster such that *τ*_*NM*_ approaches 5 minutes, the leakage of active calcineurin will also become affected. Accordingly, very slow neck plasticity (up to *τ*_*NM*_ = 30 minutes) towards narrowed and elongated morphology caused an increase in LTP magnitude, as AMPARstarP leakage reduced and the receptors remained trapped within the stimulated spine. However, smaller *τ*_*NM*_ further simultaneously started restricting calcineurin leakage. This caused stronger AMPARstarP dephosphorylation within the stimulated spine head and, hence, a decline in LTP magnitude. Similarly, slow neck plasticity (up to *τ*_*NM*_ = 10 minutes) towards widened-and-shortened morphology predominantly increased AMPARstarP leakage and caused a gradual decrease in LTP magnitude. However, further decreases in *τ*_*NM*_ caused stronger calcineurin leakage, leading to reduced AMPARstarP dephosphorylation in the stimulated spine head, and an increase in the LTP magnitude. Extremely fast seconds-timescale *τ*_*NM*_ was associated with very strong leakage of both AMPARstarP and calcineurin, which lead to net small LTP magnitude. Stronger AMPARstarP leakage with faster neck septin plasticity (decreasing *τ*_*NS*_) caused decrease in the overall LTP magnitude observed under neck morphological plasticity.

## Discussion

Using computational modelling, we studied the role of spine neck plasticity in the expression of LTP at single synapses. Our biophysical model of AMPAR trafficking and intracellular signalling included CaMKII and calcineurin dynamics in spines on an apical CA1 dendritic branch. We examined the differential contributions of plasticity in spine neck morphology and the septin7 ring, which can restrict the diffusion of membrane proteins (here AMPARs), across the base of the spine neck. Our main result is that neck morphological plasticity substantially impacts LTP magnitude. Plasticity towards narrowed and elongated neck morphology with increased neck restrictions to both membrane diffusion of AMPARs and cytosolic diffusion of calcineurin caused a large decrease in LTP magnitude. Interestingly, neck plasticity towards wider and shorter neck morphologies that are less restrictive also moderately reduced LTP magnitude, but via different mechanisms. Together these results imply that there may be an optimal spine neck morphology that maximizes LTP magnitude. In our simulations, it was the original neck configuration, which we chose to match the statistically most frequent values of neck radius and length observed in empirical data [7,21,43]. Spine neck plasticity involving changes in the neck septin7 ring complexes, with or without neck morphological plasticity, revealed that septin7 barriers at the base of the neck maximize LTP magnitude. Finally, we found that the timescale of spine neck plasticity was a key determinant of the magnitude of LTP.

Studies have found that synaptic LTP is paralleled by spine neck changes towards a wider and shorter geometry [27,28,70]. However, these studies typically focused on the electrophysiological consequences of neck plasticity, where wider and shorter neck geometries are predicted to reduce neck electrical resistance and potentially affect EPSP amplitude [27,28]. However, interpreting the data from these studies is difficult because both synaptic conductance and neck electrophysiological properties change in parallel, making it hard to dissociate their relative contributions to EPSP amplitude following plasticity. Computational modelling may prove useful for characterising these phenomena. In our study we did not model synapse or spine electrophysiology, but instead found that neck morphological plasticity can also directly impact synaptic AMPAR count, a key determinant of synaptic conductance, during LTP by modulating the AMPAR trafficking through biochemical signalling alone. Future experimental and computational studies can consider the joint, interactive effects of spine neck plasticity on electrophysiological and biochemical signalling underlying LTP.

Interestingly, Tonnesen et al. [28] found that spine neck morphological plasticity was typically paralleled by changes in spine head size. These changes seemed to be dynamically co-ordinated at the single-spine level, despite the fact that there is typically little or no correlation observed between these properties at the population-level in ultrastructural data [7,21,23,43,44]. Spine head plasticity is important here because head size is also believed to affect the time constant of both cytoplasmic and membrane diffusion of molecules out of the spine head: larger heads should tend to trap molecules for longer [14,16,62]. In our study we did not systematically explore the effects of spine head plasticity on AMPAR dynamics and LTP, but instead directly replicated in our model the dynamics of spine head plasticity (Figure 2B.i) from averaged fluorescence imaging data from Patterson et al, 2010 [41]. It would be interesting for future studies to explore the interactions between spine head and neck plasticity.

One important conclusion from our study is that spine neck coupling for both membrane and cytosolic diffusion cannot be simultaneously preserved when changing neck morphology alone, as membrane diffusion rate is directly proportional to the neck radius whereas cytosolic diffusion rate varies with the square of the neck radius (Figure 7). For example, doubling the diameter of the spine neck would lead to a two-fold decrease in the membrane diffusion time constant, but a four-fold decrease in cytoplasmic diffusion time constant. In contrast, changes to spine neck length should affect time constants for cytoplasmic vs membrane diffusion equally. These observations have implications beyond AMPA dynamics, and apply to all cytoplasmic and membrane-diffusing molecules [71]. In addition to this differential but interlinked relationship between neck morphology and membrane vs cytoplasmic molecular diffusion, there may also be several additional mechanisms that can independently regulate diffusion via one or the other routes. In our study we considered only the septin7 ring, which can form a barrier at the base of the spine neck to selectively modulate membrane diffusion [24]. Similarly, synaptopodin in the spine neck might participate in regulating membrane diffusion via the F-actin cytoskeleton [72]. Conversely, other mechanisms such as actin filaments [73] or the spine neck apparatus [74] may regulate cytoplasmic diffusion. Whether and how all these various mechanisms are jointly controlled by synaptic signalling pathways remains an important open question.

**Figure 7.**
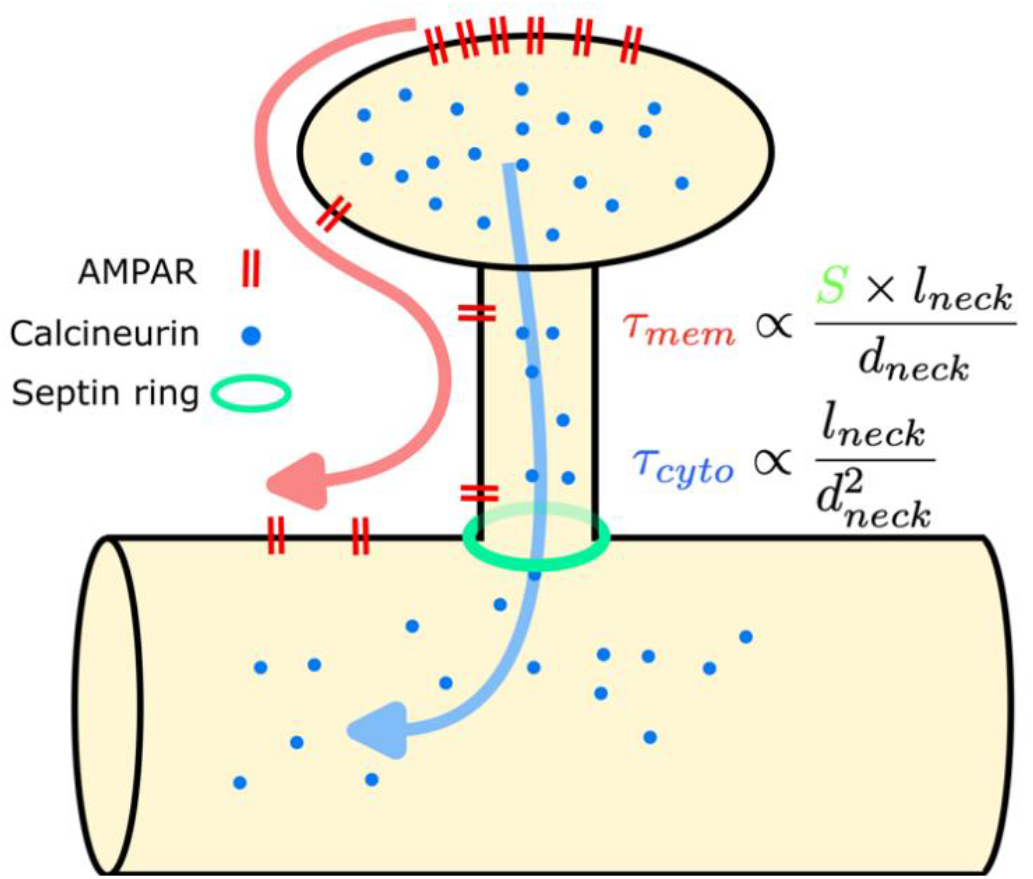
Spine neck morphological and molecular properties control the balance between AMPAR and calcineurin diffusion from the spine. AMPARs (red) diffuse between the spine head and the dendritic shaft via the spine neck membrane. In contrast, calcineurin (blue) diffuses between the spine head and the dendritic shaft via the spine neck cytoplasm. The time constants of spine-dendrite exchange for the two diffusion processes (equations on right) have differing relationship with spine neck properties: 1) membrane diffusion scale proportional to neck diameter, *d*_*neck*_, while cytoplasmic diffusion scales with the square of neck diameter; 2) membrane, but not cytoplasmic, diffusion can be impeded by the septin ring barrier, *S*, (green hoop) at the base of the spine neck.

This work generates several experimentally testable predictions: 1) blocking exocytosis will result in a fast but short-lasting LTP, while blocking AMPA phosphorylation will result in a slower-rising but long-lasting LTP (Figure 3A). 2) For a spine with typical neck dimensions, blocking neck structural plasticity will enhance LTP (Figure 5B). 3) During LTP-stimulation, spine neck narrowing or lengthening causes increased calcineurin-based dephosphorylation of AMPARs, whereas spine neck shortening or widening causes enhanced phosphorylated AMPAR escape. 4) Removing the spine neck septin barrier will reduce LTP magnitude, by allowing AMPARs to diffuse faster from the spine head (Figure 5A). 5) Either slowing down or speeding up the dynamics of neck plasticity affects LTP magnitude (Figure 6).

Following LTP stimulation in the model, we observed a moderate accumulation of phosphorylated AMPARs and the phosphatase calcineurin in the dendritic shaft (Figures 5C and 5D). Experimentally, these two molecules are reported to diffuse into neighbouring spines [14,47,59,62] where they could potentially either directly lead to plastic changes in the strengths of synapses, or modulate their propensity for plasticity. Here we simulated only a single-spine stimulation protocol, but if multiple neighbouring spines were stimulated simultaneously, as likely occurs *in vivo*, then these spillover signals could accumulate to induce substantial heterosynaptic plasticity effects [75-78]. Data-driven computational modelling of these processes may be useful for dissecting out the contributions of these various interacting spatiotemporal processes in dendrites during learning.

## Acknowledgements

This work was funded by grants from the Leverhulme Trust (RPG-2019-229) and Medical Research Council (MR/S026630/1) to COD. A subset of the simulations was run on the University of Bristol’s Blue Crystal Phase 4 high-performance computing service.

## Methods

### AMPAR Trafficking dynamics

We designed a compartment-based reaction-diffusion framework for the modeling of AMPAR trafficking dynamics. It consisted of a system of coupled first-order ordinary differential equations (ODEs). Given the length scales of the membrane compartments we chose (1 *μm*), in real neurons the diffusion of AMPARs would be expected to quickly homogenize the receptor distribution in a compartment within a few seconds. Accordingly, we assumed that a mass-model approach of homogenous receptor density in a compartment during inter-compartment receptor exchange could be implemented for studying the AMPAR trafficking dynamics at the minutes timescale [33, 34].

AMPARs were assumed to always exist in complexation with the stargazin proteins [45]. The AMPAR-stargazin population was comprised of two distinct species: unphosphorylated (AMPARstar) and phosphorylated (AMPARstarP) complexes. For brevity of notation below, we denote the AMPARstar as ‘U’ for ‘Unphosphorylated’ whereas AMPARstarP complexes as ‘P’ for ‘Phosphorylated’. The densities of the individual AMPAR species (*R*) were denoted as *R*^*i*^, where *i* ∈ {*U, P*} denoted the phosphorylation tag. Similarly, the count of individual AMPAR species in the intracellular pool is denoted as *IR*^*i*^, where *i* ∈ {*U, P*}.

In the PSD region of the spine head, AMPARs could either be freely diffusing with diffusion coefficient *D*_*PSD*_ or be bound to the PSD95 slot proteins and hence unable to diffuse. This was based on high-resolution fluorescence-imaging and single particle tracking studies that show a mixture of diffusing and spatially-frozen PSD95-bound receptors in the PSD [79-81]. The dynamics of the freely-diffusing 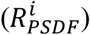 and PSD95-bound 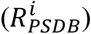 AMPAR species in the PSD were given by,

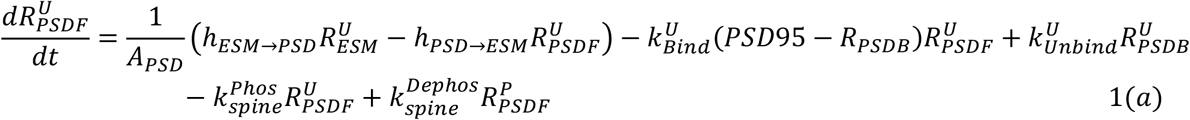

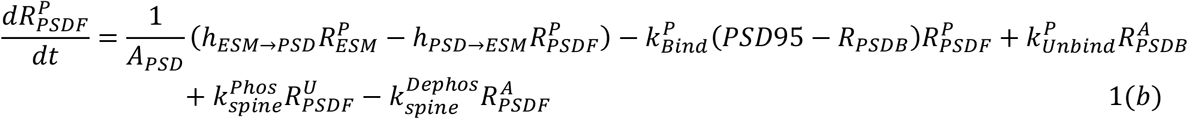

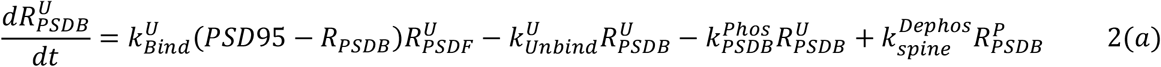

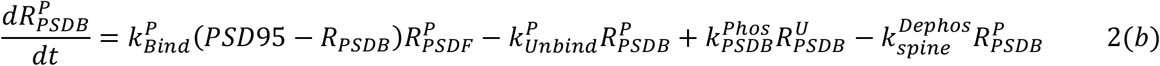

The first right-hand side terms of Equations 1(a) and 1(b) describe the exchange of freely-diffusing individual AMPAR species between the PSD and the ESM. 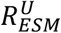 and 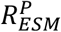 denote the densities of AMPARstar and AMPARstarP receptor species, respectively, in the spine ESM. The parameters *h* represent the rate of hopping (units, *μm*^2^. *s*^−1^) of the AMPARs between the compartments mentioned in the subscript, with an arrow depicting the direction of hopping. Since the PSD is assumed as a flat circular patch in the spine membrane, the hopping parameters are defined as,

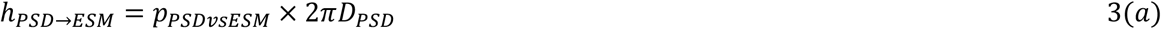

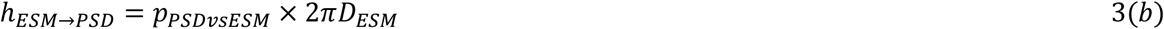

*p*_*PSDvsESM*_ denotes the bidirectionally symmetric permeability of the boundary between PSD and ESM to the receptor flux. It took care of the fact that the crowded presence of several transmembrane proteins which coral/fence the PSD boundary [82,83] or physical barriers at the boundary arising from lipid structures [84] may impede a smooth entry or exit of the AMPARs, and only narrow escapes are possible. Here, we kept *p*_*PSDvsESM*_ = 1. The AMPARstar and AMPARstarP species shared identical hopping parameters here as well as in the remaining membrane compartments, as phosphorylation-dependent changes in the AMPAR diffusion coefficient are not yet known.

The second and third terms on the right-hand side of Equations 1(a) and 1(b) as well as of Equations 2(a) and 2(b) describe the binding-unbinding process of the respective AMPAR species with PSD95. Here, PSD95 denotes the density of PSD95 slot proteins and 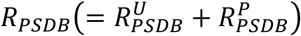 represents the density of total bound receptors. Therefore, (*PSD*95 − *R*_*PSDB*_) shows the density of free PSD95 slot proteins available for binding. 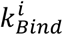 and 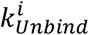, *i* ∈ {*U, P*}, represent the binding and unbinding rate constants of the individual AMPAR species. Here we assumed 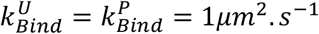 always. We rather expressed the differential binding of AMPAR species to the PSD95 in terms of the ratios of their unbinding to binding rate constants, as

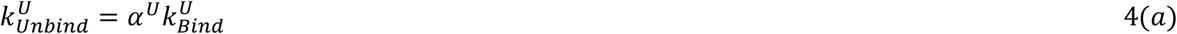

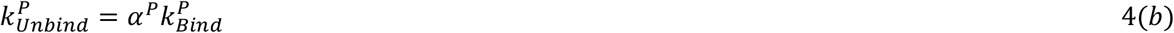

*α*^*U*^ and *α*^*P*^ are the ratios of unbinding to binding rate constants of AMPARstar and AMPARstarP species. Stronger binding of AMPARstarP to PSD95 in comparison to that of AMPARstar was conceived as *α*^*P*^ ≪ *α*^*U*^.

The last two terms on the right-hand side of Equations 1(a) and 1(b) describe the phosphorylation-dephosphorylation of the free AMPARs and the last two terms on the right-hand side of Equations 2(a) and 2(b) describe the same for bound AMPARs. 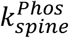 and 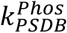 represent the different phosphorylation rate constants of the freely-diffusing and PSD95-bound AMPARstar, respectively, in the PSD region. Here, 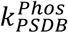 is always greater than 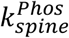, as CaMKII preferentially binds PSD95 [85,86] and are frequently located close to the receptor binding slots. However, both free and bound AMPARstarP species shared an identical dephosphorylation rate constant, 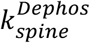. Calcineurin is relatively more homogeneous in the spine volume and, hence, all membrane receptors in the spine head are exposed to an identical calcineurin concentration.

The dynamics of the individual AMPAR species in the spine ESM are given by,

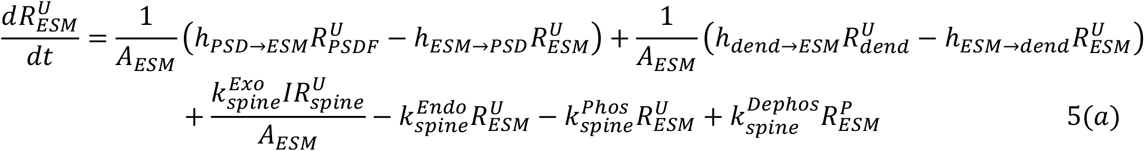

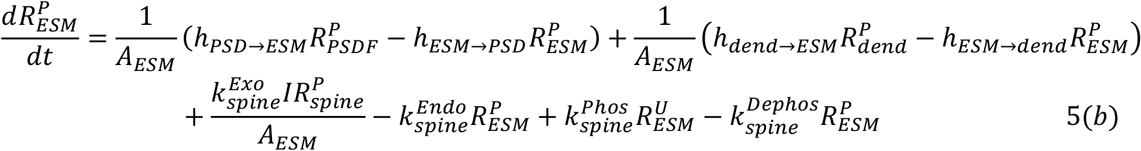

The first terms on the right-hand side of Equations 5(a) and 5(b) describe the exchange of AMPARs between PSD and ESM whereas the second terms describe that between the spine head and the dendrite shaft. 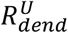 and 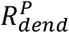 denote the densities of AMPARstar and AMPARstarP species in the membrane of dendritic shaft compartment receiving the spine neck. The hopping factor *h*_*ESM*→*dend*_ is given by,

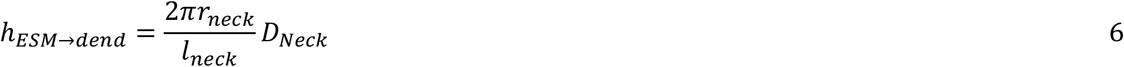

and *h*_*dend*→*ESM*_ = *h*_*ESM*→*dend*_ by symmetry. *D*_*Neck*_ represents the effective diffusion coefficient of AMPARs in the spine neck region. The third and fourth terms on the right-hand side of Equations 5(a) and 5(b) describe the receptor exocytosis and endocytosis. 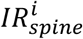, *i* ∈ {*U, P*} represents the intracellular receptor pools of the AMPARstar and AMPARstarP in the spine head. The parameters 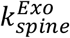 and 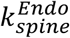 denote the rate constants of exo- and endo-cytosis (units, *s*^−1^) of receptors and were considered identical for both the AMPAR species. The last two terms on the right-hand side of Equations 5(a) and 5(b) describe the phosphorylation-dephosphorylation of the AMPARs in the spine ESM.

The dynamics of the intracellular receptor pools are given by,

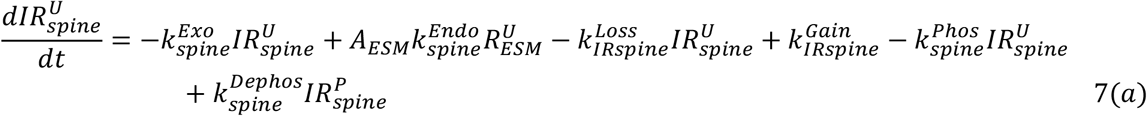

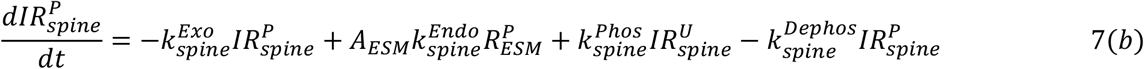

Here, the first two terms on the right-hand side of Equations 7(a) and 7(b) are associated with receptor exocytosis and endocytosis. The third term of Equation 7(a) describes the loss of receptor pool through lysosomal degradation [50] and active retrograde vesicular transport of receptors from dendritic spines [51,52]. Both the processes occur at minutes timescale and can be dealt collectively with a single rate constant. 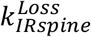 is the rate constant of receptor loss in the IR pool of spine head (units, *s*^−1^). The fourth term on the right-hand side of Equation 7(a) describes the constant rate of receptor gain through local synthesis of new receptors [50] as well as active anterograde transport of receptors to spines [51,52]. 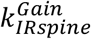 is the rate constant of receptor gain in the IR pool of spine head (units, *s*^−1^). Here too, the receptor synthesis and anterograde transport occur at minute timescale and, hence, can be grouped together. The last two terms on the right-hand side of Equations 7(a) and 7(b) describe the phosphorylation and dephosphorylation of the intracellular pools. We assumed that the phosphorylated receptor pool did not undergo direct loss and gain through synthesis/degradation and active motor endosomal transports. It is only enriched through phosphorylation of the unphosphorylated receptor pool and depleted through dephosphorylation.

The dynamics of membrane receptors in the *kth* dendritic shaft compartment bearing a dendritic spine are given by,

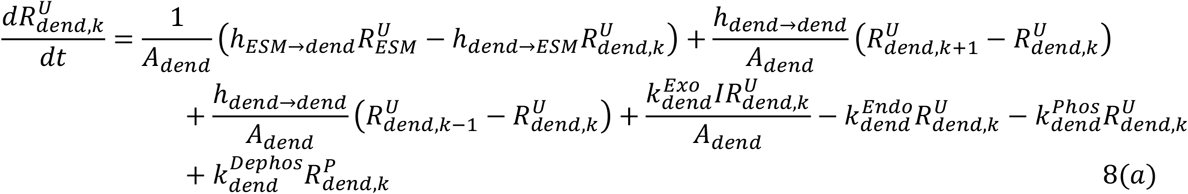

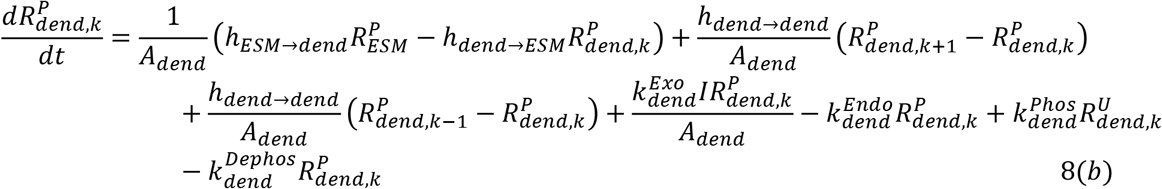

The first terms on the right-hand side of Equations 8(a) and 8(b) represent the exchange of individual AMPAR species between the spine head and the dendritic shaft compartment. The second and third terms on the right-hand side of Equations 8(a) and 8(b) represent the exchange of receptors between the *kth* compartment and the immediate neighbouring compartments, *k* + 1 and *k* − 1, of the dendritic shaft. The hopping parameter *h*_*dend*→*dend*_ denotes the rate of receptor hopping amongst the dendritic shaft compartments and is given by,

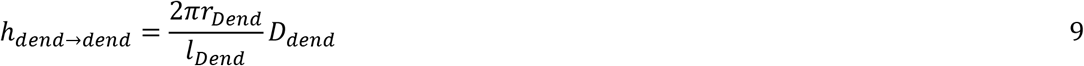

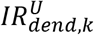 and 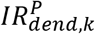 denote the size of the IR pools of individual AMPAR species in the *kth* shaft compartment. 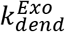 and 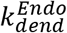 are the exocytosis and endocytosis rate constants (units, *s*^−1^). The last terms of both the equations denote the phosphorylation and dephosphorylation dynamics with the respective rate constants 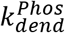 and 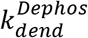. The dynamics of the intracellular receptor pools are given by,

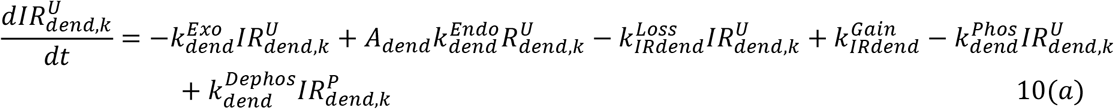

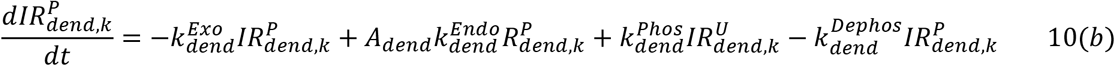

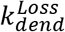 and 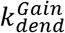 denote the rate constants of the loss and gain of intracellular receptors through mechanisms discussed above for the spine intracellular receptor pools. The last pair of terms on the right-hand side of Equations 10(a) and 10(b) represent the phosphorylation and dephosphorylation of intracellular receptors.

The boundary conditions of the dynamical model are given as,

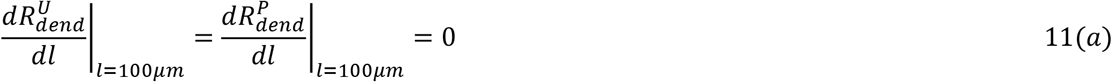

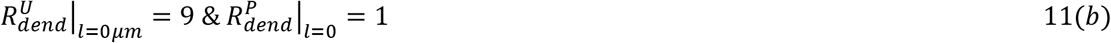

*l* = 0*μm* denotes the origin of the secondary dendritic branch from its parent dendrite whereas *l* = 100*μm* denotes the distal tip of the branch. Equation 11(a) describes the perfectly reflective or no-passage boundary condition for diffusing membrane receptors at the shaft’s distal tip. Equation 11(b) describes the time-invariant steady baseline densities of the individual receptor species in the dendritic shaft membrane at the point of branch origin.

### Steady State Analysis and Parameter Estimation for Baseline Condition

The geometrical features of all the dendritic spines on the uniformly thick dendritic shaft were identical in the initial unstimulated baseline condition of the model. Further, we considered perfectly homogeneous distributions of the AMPARstar and AMPARstarP species in the membrane as well as intracellular compartments as well as of the active CaMKII and calcineurin levels across all the dendritic spines and shaft. We also considered identical rate kinetics of the different AMPAR trafficking and signalling processes across the dendritic spines and shaft under the baseline condition. Table 1 lists the geometric parameters, the densities/counts of the AMPAR species, and the concentration of active calcineurin under the baseline condition.

**Table 1.** Geometric parameters of the dendritic branch and densities/counts of AMPARs and calcineurin under the initial baseline condition of the model. ‘Tot.’ refers to the total density of AMPAR including the AMPARstar and AMPARstarP species. The ‘*’ in the superscripts of the notations for AMPAR densities refers to the steady state densities. ‘U’ and ‘P’ in the superscripts respectively refer to the unphosphorylated AMPARstar and phosphorylated AMPARstarP species. *i* ∈ {*PSDF, ESM, neck, dend*} and *j* ∈ {*spine, dend*}.

| Parameters | Values | Ref. |
| --- | --- | --- |
| Spine head surface area, $A_{spine}$ | $1.1 \mu m^2$ | 7,21,43 |
| Spine head volume, $V_{spine}$ | $0.1 \mu m^3$ | 7,21,43 |
| Area of ESM, $A_{ESM}$ | $1 \mu m^2$ | 21,43,44 |
| Area of PSD, $A_{PSD}$ | $0.1 \mu m^2$ | 21,43,44 |
| Spine neck length, $l_{neck}$ | $0.3 \mu m$ | 7,21,43 |
| Spine neck radius, $r_{neck}$ | $0.0562 \mu m$ | 7,21,43 |
| Dendritic shaft radius, $r_{dend}$ | $0.5 \mu m$ | 42 |
| PSD95 density in PSD, $PSD95$ | $1000 \mu m^{-2}$ | 46,57,58 |
| Freely diffusing tot. AMPAR density in PSD, $R_{PSDF}^*$ | $100 \mu m^{-2}$ | This paper |
| PSD95-bound tot. AMPAR density in PSD, $R_{PSDB}^*$ | $250 \mu m^{-2}$ | 46,57 |
| Tot. AMPAR density in ESM, $R_{ESM}^*$ | $10 \mu m^{-2}$ | 55,56 |
| Tot. AMPAR density in spine neck membrane, $R_{neck}^*$ | $10 \mu m^{-2}$ | 55,56 |
| Tot. AMPAR density in dendritic shaft membrane, $R_{dend}^*$ | $10 \mu m^{-2}$ | 55,56 |
| Intracellular tot. AMPAR pool of spine head, $IR_{spine}^*$ | 100 | This paper |
| Intracellular tot. AMPAR pool of dendritic shaft compartment, $IR_{dend}^*$ | 10 | This paper |
| Ratio of membrane AMPARstar:AMPARstarP, $R_i^{U,*} : R_i^{P,*}$ | 9:1 | This paper |
| Ratio of intracellular AMPARstar:AMPARstarP, $IR_j^{U,*} : IR_j^{P,*}$ | 9:1 | This paper |
| Ratio of PSD95-bound AMPARstar:AMPARstarP, $R_{PSDB}^{U,*} : R_{PSDB}^{P,*}$ | 1:9 | This paper |
| Active calcineurin concentration in entire dendritic branch, $[Can]$ | $0.1 \mu M$ | 36,39 |

Since we had already set the desired baseline densities of the individual AMPAR species in the different membrane compartments and the receptor counts in the IR pools, we could compute the unknown kinetic parameters using steady analysis of Equations 1-10. We first assumed that the phosphorylation-dephosphorylation flux (the last pair of terms on the right-hand side of Equations 1-10) are mutually balanced in all the compartments as,

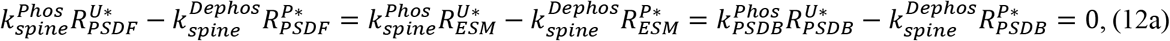

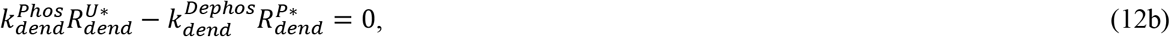

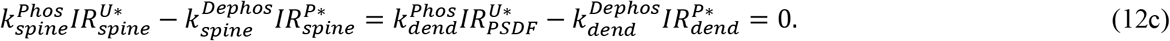

The asterisk in the superscript represents the steady state density/counts of the individual AMPAR species in the various compartments. This implied that, under the baseline condition, the ratio of the phosphorylation and dephosphorylation rate constants in a compartment is identical to the ratio of the steady state AMPARstarP to AMPARstar species in that compartment. The ratio of steady state densities of AMPARstarP to AMPARstar species in all the membrane compartments and that of the steady state counts in the intracellular pools were 1:9. The ratio was reversed (9:1) for the PSD95-bound AMPAR population. In this way, the phosphorylation and dephosphorylation rate constants under the baseline condition were constrained with each other as,

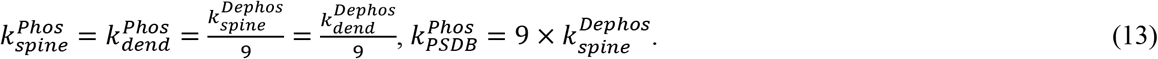

Note that we had assumed a homogenous distribution of active CaMKII and calcineurin across the dendritic spines and shaft. The phosphorylation rate constant was higher only for the bound AMPARs as the active CaMKII intracellularly accumulated around the PSD95 proteins within the PSD region [85,86]. Therefore, knowing the value of 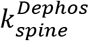 would inform us about the baseline values of the phosphorylation rate constants.

As described previously, we modelled the differential binding of AMPARstar and AMPARstarP to the PSD95 slot proteins in terms of the different ratios of the unbinding to binding rate constants, *α*^*U*^ and *α*^*P*^, respectively. The steady state analysis of Equations 2a and 2b provided a constrained relationship between the *α*^*U*^ and *α*^*P*^ as,

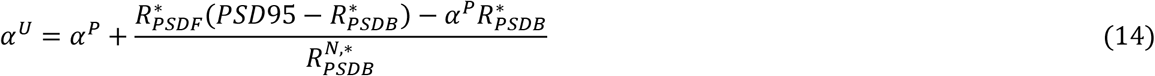

Here, 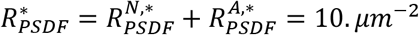, previously fixed in the model. Therefore, knowing *α*^*P*^ would spontaneously yield *α*^*U*^. Eventually, this led to the computation of unbinding rate constants, 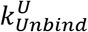 and 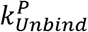, given the identical binding rate constants 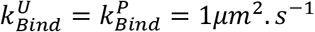 [87] we assumed in the model.

The hopping parameters for the diffusion of AMPARs amongst the membrane compartments were easily computed (Equations 3a, 3b, 6 and 9), given the fixed geometric features of the compartments and the empirical estimates of the receptor diffusion coefficients in the membrane compartments. The remaining eight trafficking parameters to be estimated were the exocytosis-endocytosis rate constants and the loss-gain rate constants for the intracellular receptor pool dynamics, each for the spine head and the dendritic shaft compartments. However, steady-state analysis reduced it to four parameters, where the endocytosis and the gain rate constants were constrained to the exocytosis and the loss rate constants, respectively, in the spine head and the dendritic shaft. The endocytosis rate constant was constrained to the exocytosis rate constant as,

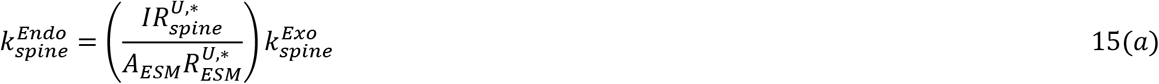

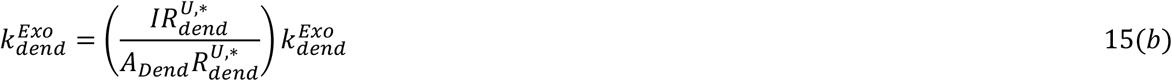

Similarly, the baseline AMPAR gain rate constant in the intracellular receptor pool was constrained to the loss rate constant as,

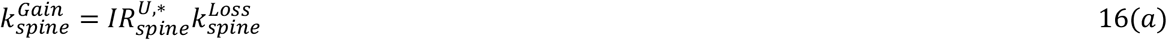

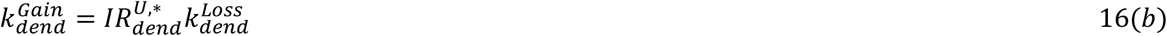

Altogether, we had six parameters to be estimated: 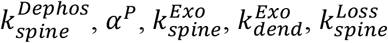, and 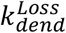. For the parameter estimation, we used the imaging data from Patterson et al. study [41] on the fluorescence recovery of surface AMPARs after photobleaching (FRAP) of the apical dendritic branch under the unstimulated resting condition. First, we formulated a FRAP version of our original model. Since only the surface AMPARs were photobleached in the experimental study, we considered two identical copies of the original system of coupled ODEs (Equations 1a, 1b, 2a, 2b, 5a, 5b, 8a, and 8b) for the membrane AMPAR dynamics of the unbleached and photobleached surface AMPARs. The dynamics of intracellular pools (Equations 7a, 7b, 10a, 10b) were unaffected and contained only unbleached AMPARs. Therefore, the exocytosis rate constants for the photobleached surface AMPARs were identically zero in the spine as well as the dendritic shaft compartments. Finally, to estimate the unknown parameters, we performed an automatic optimization of the FRAP version of the model over the six-dimensional parameter space to fit the empirical FRAP data.

Estimation yielded *α*^*P*^ = 33.3 *μm*^−2^ and 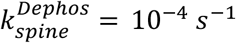. Using Equation 14, we obtained *α*^*U*^ = 2700 *μm*^−2^. Using Equation 13, we obtained 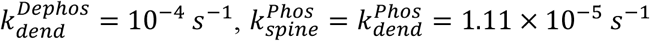, and 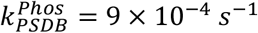. Notably, such low levels of phosphorylation and dephosphorylation rate constants in the initial baseline condition signify steady but extremely slow signalling processes in the unstimulated dendritic branch. The estimated value of baseline exocytosis rate constant in spine head was 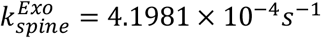. Patterson et al. [39] showed that each quantal event of vesicle fusion led to a 50% rise in the total surface fluorescence of spine head during resting condition. In our model, since the total count of membrane AMPARs in the spine head was 45 and an intracellular pool of 100 AMPARs in the baseline condition, the estimated exocytosis rate constant is equivalent to 0.1120 *events. min*^−1^, which is almost identical to the experimentally observed rate of 0.11 ± 0.05 *events. min*^−1^ [39]. The estimated exocytosis rate constant of AMPARs in the dendritic shaft turned out 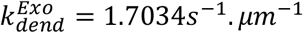, which is equivalent to 0.6506 *events. min*^−1.^ *μm*^−1^ (experimental estimate: 0.034 *events. min*^−1.^ *μm*^−1^ [39]). Using steady-state analysis (Equations 15a and 15b), the baseline endocytosis rate constants in the spine head and the dendritic shaft turned out 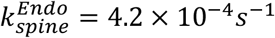 and 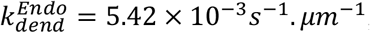, respectively. The estimated loss and gain rate constants (Equation 16a) for the AMPAR intracellular pool in the spine head were 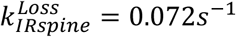 and 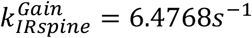 (for dendritic shaft: 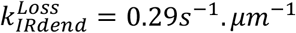 and 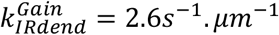, respectively). Table 2 lists the values of all the kinetic parameters in the model.

**Table 2.**
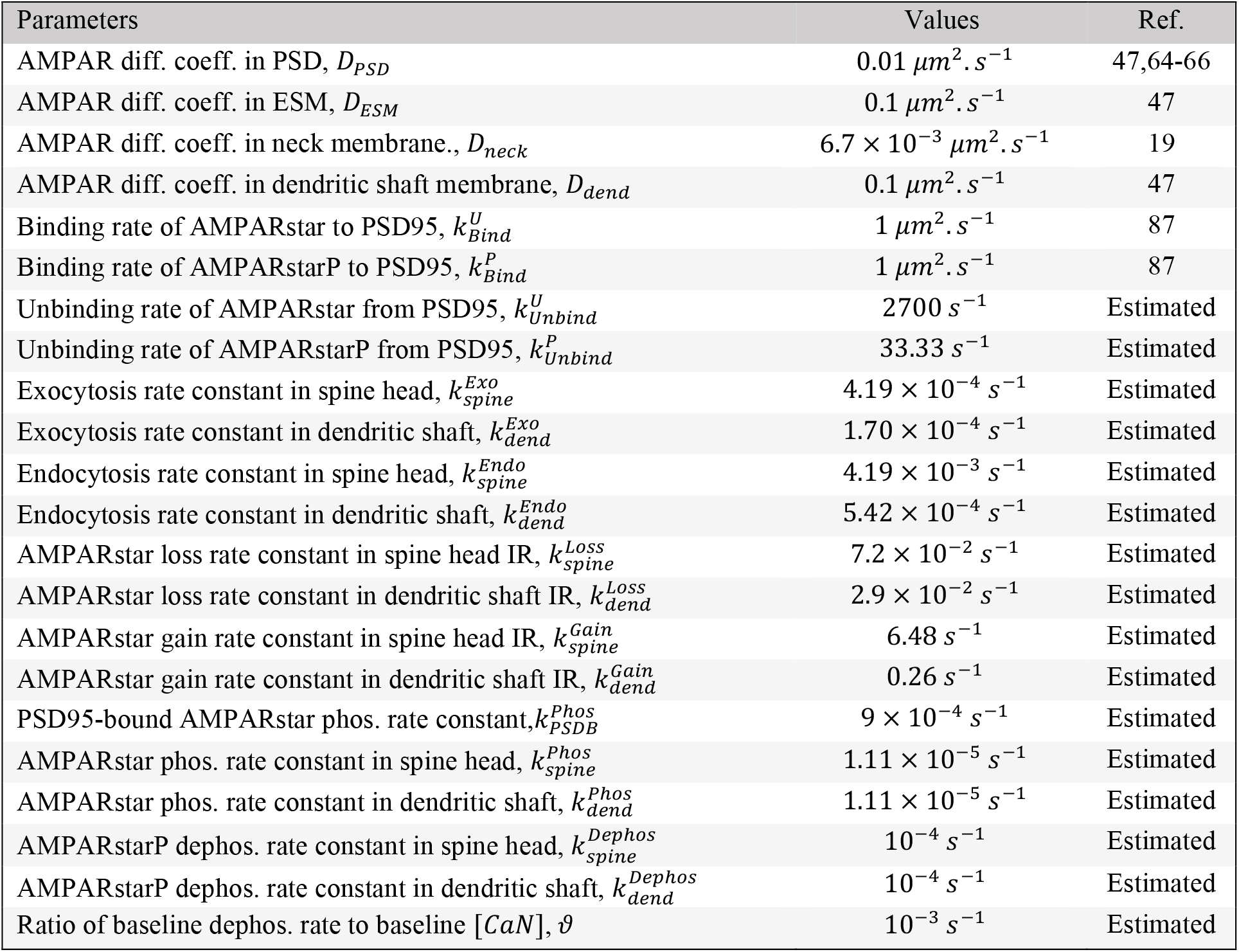
Kinetic parameters of the model under the baseline unstimulated condition. The parameters which were estimated in the present model are mentioned as ‘Estimated’ in short. ‘Diff. coeff.’ stands for diffusion coefficient. IR refers to the intracellular receptor pool. ‘Phos.’ and ‘dephos.’ stand for phosphorylation and dephosphorylation, respectively. [*CaN*] denotes the concentration of active calcineurin.

### Signaling Dynamics of CaMKII and Calcineurin

LTP-inducing glutamatergic stimulation of a dendritic spine causes activation of the synaptic NMDA receptors, which leads to local influx of Ca^2+^ in the stimulated spine head. Ca^2+^ activates cytosolic calcium-sensor proteins calmodulin, which eventually leads to activation of CaMKII and calcineurin enzymes. This entire signaling process involves multiple nonlinear reaction steps [35,38,59]. We skipped the large and intricate dynamical framework of the signaling cascade itself. Instead, we directly varied the levels of active CaMKII and calcineurin in the stimulated spine head in accordance with the profiles observed in the previous experimental studies [60,61] as well as the computational studies involving detailed modelling of the signaling cascade [35,38]. In our model, the concentrations of active CaMKII and calcineurin enzymes drove the phosphorylation and dephosphorylation rate constants, respectively, of the AMPARs in the membrane compartments as well as the intracellular pools. Experimental studies have shown that activated CaMKII remains localized in the stimulated spine head [14,60-63]. Since active CaMKII lacked any spatial dynamics, we directly varied the phosphorylation rate constants in the stimulated spine head to capture the effect of locally elevated active CaMKII during spine stimulation. However, activated calcineurin exhibits significant spillage out of the stimulated spine head and spread into the neighboring dendritic stretch [14,62]. Therefore, we explicitly modelled calcineurin as a diffusible enzyme and considered its spatiotemporal dynamics.

Fluorescence imaging studies have shown that the concentration of active CaMKII in the spine head rapidly rises within a few seconds of glutamate application and attains an elevated plateau over the duration of spine stimulation [60,61,63]. However, at the end of glutamate application, the level of active CaMKII rapidly decays to the background level within around one minute post-stimulation. Computational studies on the detailed signaling dynamics [35,38] have also shown a similar profile of variation in the level of active CaMKII and calcineurin in the stimulated spine head. Further, like CaMKII, the concentration of active calcineurin also decays to the background level with a minute or two post-stimulation [14,62]. Accordingly, we approximated these empirical temporal profiles with a step-increase in the levels of the active CaMKII and calcineurin in the stimulated spine head at the beginning of glutamate application, followed by a sustained or clamped elevated levels over a minute long glutamate application, and an exponential decay to the background levels with different time constants, *τ*_*CaM*_ and *τ*_*Can*_ respectively, post-stimulation. To model CaMKII, we performed the approximated temporal variation in the phosphorylation rate constants, 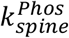 and 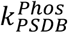, in the stimulated spine head. In the case of calcineurin, the active enzyme not only underwent self-decay but also diffusion out of the spine and along the dendritic stretch with a diffusion coefficient *D*_*Can*_.

As estimated above, the baseline phosphorylation rate constants in the spine head were 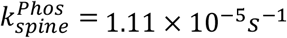 and 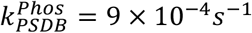, which is specifically for PSD95-bound AMPAR pool. The temporal profile of variation in the phosphorylation rate constants in the stimulated spine head is given by,

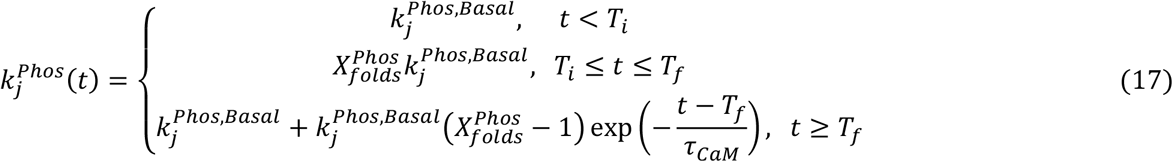

, where *j* ∈ {*spine, PSDB*}, 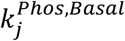 is the baseline value of the phosphorylation rate constant, and 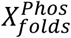 is the folds increase in the phosphorylation rate constant. Note that we made identical folds increase in both the kinds of phosphorylation rate constant. *T*_*i*_ and *T*_*f*_ are the time points of beginning and end of the glutamatergic stimulation. Since CaMKII did not diffuse out of the stimulated spine, the phosphorylation rate constant in the dendrite remained invariant from its basal value, which was identical to the baseline value of 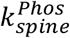.

The baseline dephosphorylation rate constants in the spine head as well as the dendritic shaft were 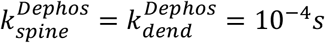. Since we modelled the calcineurin enzyme explicitly, a baseline concentration of active calcineurin, [*CaN*]^*Basal*^ = 0.1*μM*, was chosen homogenously distributed across all the dendritic spines and shaft. The spatiotemporal dynamics of the concentration of active calcineurin, in the dendritic branch was given by,

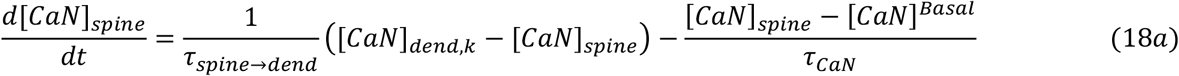

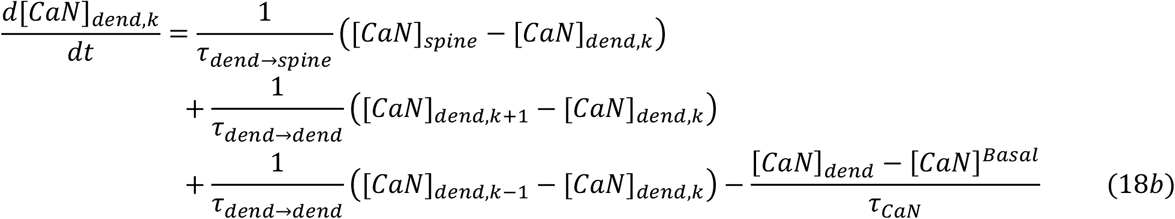

[*CaN*]_*spine*_, [*CaN*]_*dend,k*_, [*CaN*]_*dend,k*+1_ and [*CaN*]_*dend,k*−1_ denote the concentration of active calcineurin in the dendritic spine, in the *kth* dendritic shaft compartment bearing dendritic spine, in the neighboring one shaft compartment upwards and in the one shaft compartment downwards, respectively. The first terms on the right-hand side of Equations 18(*a*) and 18(*b*) describe the exchange of active calcineurin between the spine head and the dendritic shaft compartment. The second and third terms of Equation 18(*b*) describe the same amongst the neighbouring shaft compartments. The last terms of both the equations describe inactivation of active calcineurin to its basal value in the spine head and the shaft compartment. The *τ*_*spine*→*dend*_ and *τ*_*dend*→*spine*_ are the spine neck coupling time constant for the intracellular volume diffusion of calcineurin from spine head to the dendritic shaft compartment and are given by,

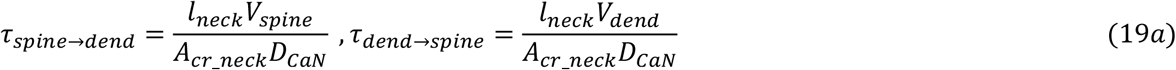

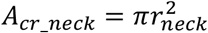, denotes the cross-sectional area of the spine neck. *V*_*spine*_ and *V*_*dend*_ denote the volumes of the dendritic spine head and shaft compartment, respectively. Similarly, *τ*_*dend*→*dend*_ was given by,

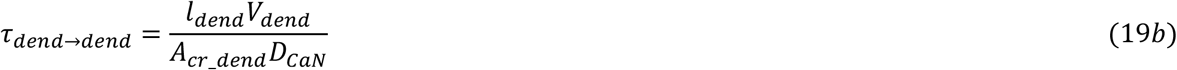

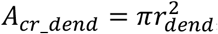, denotes the cross-sectional area of the dendritic shaft. During the glutamatergic stimulation from *T*_*i*_ to *T*_*f*_, the level of active calcineurin within the stimulated spine head was clamped to 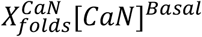, where 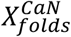 is the folds increase in the active calcineurin level. Based on the concentration of active calcineurin in a compartment, the local dephosphorylation rate constant varied as 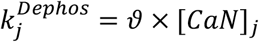, where *j* ∈ {*spine, dend*} and *ϑ* = 10^−3^, which is the ratio of the baseline dephosphorylation rate constant to the baseline concentration of active calcineurin.

### Spine stimulation protocol

For the single spine stimulation, we chose the middle-most spine (located at 50um from the branch origin, Figure 2B) in our model. We implemented the LTP-inducing spine stimulation in three parts:

1. To incorporate the dynamics of structural plasticity of the stimulated spine head, we directly increased the size of the spine head in a manner identical to the temporal profile of spine head enlargement observed by Patterson et al. [41, see Figure 1D]. However, it was only the size of the spine ESM (*A*_*ESM*_) which we changed in this manner (Figure 2B.i). The size of the PSD (*A*_*PSD*_) and the number of PSD95 slot proteins were not altered, as both have been experimentally observed [67,68] to exhibit insignificant changes during the first 30-50 minutes post-stimulation. The time span of our simulation was 30 minutes post-stimulation, which typically belongs to the early phase of LTP [68].
2. We step-increased the exocytosis rates of the AMPAR species in the stimulated spine by 26-folds in the first 50 seconds duration of the glutamate uncaging, followed by a 6-fold increase for next 10 seconds, and they set to the baseline rate eventually (Figure 2B.ii). This was in accordance with the folds-increase in quantal fusion events reported by Patterson et al. [41, Figure 4A]. Similarly, the exocytosis rates of the AMPAR species within a 5*μm* stretch of the dendritic shaft underneath the stimulated spine was step-increased for 50 seconds duration of the glutamate uncaging [41, Figure 4B]. The amount of increase depended on the distance away from the stimulated spine (Figure 2B.v) and was identically adopted from the folds-increase experimentally reported by Patterson et al. [41, Figure 4C]. The endocytosis rate constants, and the AMPAR loss and gain rate constants of the intracellular pools, were unaffected in both the stimulated spine as well as the entire dendritic shaft.
3. At the instance of glutamate uncaging, we step-increased and clamped the phosphorylation rate constants,, 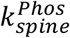 and 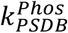, (Figure 2B.iii showing 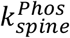) and the concentration of active Calcineurin, [*CaN*], (Figure 2B.iv) for the entire 50 seconds duration of glutamate uncaging. As mentioned above, we made a step increase of 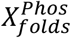 in the phosphorylation rates (Equation 17) and a step increase of 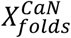 in the active calcineurin concentration in the stimulated spine head. Following the 50 seconds duration of clamped increase, the phosphorylation rate constants was allowed to exponentially decay with the time constant, *τ*_*CaM*_, to the baseline level. While the calcineurin level in the stimulated spine head was clamped, the spatiotemporal dynamics of active calcineurin occurred, according to Equations 18a and 18b. After the 50 seconds duration, the clamp was released and the spatiotemporal dynamics continued. We manually adjusted the parameters, 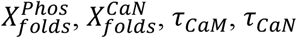, and *D*_*CaN*_, such that the temporal profile of change in the total surface AMPAR fluorescence in the stimulated spine head fit well with that reported by Patterson et al. [41, Figure 1C, SEP-GluA1 profile]. We obtained 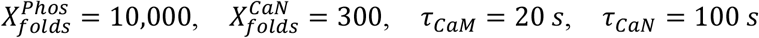, and *D*_*CaN*_ = 0.106 *μm*^2^. *s*^−1^.

### Numerical Simulation

The scripts for the numerical simulation of AMPAR trafficking and intracellular signaling dynamics were written in MATLAB R2019a (The MathWorks) and ode15s solver was used for the numerical integration of the differential equations. The scripts will be made available on request. For the parameter estimation under the resting unstimulated condition, the model was fitted to the FRAP data from Patterson et al. [41, Supplementary Fiigure 2] using the MATLAB inbuilt solver *fmincon* for nonlinear least square optimization and the default interior-point algorithm.

